# A mechanistic framework for photon-to-carbon efficiency: coupling photosynthetic energy supply and light respiration in microalgae

**DOI:** 10.64898/2026.05.07.723413

**Authors:** Guillaume Cogne, Jack Legrand, Claude-Gilles Dussap

## Abstract

Photon-to-biomass conversion efficiency in oxygenic phototrophs can vary widely under controlled cultivation, yet its mechanistic origin remains difficult to resolve at the system level. Here, we combine a quantified description of light attenuation and primary photochemistry with a thermodynamically constrained metabolic framework to relate reactor-scale light forcing to intracellular energetics in *Chlamydomonas reinhardtii* chemostats.

In a near-optimal regime, the specific growth rate scales linearly with an effective photochemical input, ϕ_γ_, defined as the rate of photochemically productive photon absorption. This identifies a physiologically relevant forcing variable at the timescale of growth and an invariant biomass yield on productive photons. Imposing this experimentally grounded constraint reveals a compatible effective energetic coupling within photosynthetic electron transport and strongly restricts the admissible photosynthesis–respiration balance.

A two-dimensional growth map in the space of photochemical forcing and respiratory activity reveals a respiration-limited region and a dissipative regime in which excessive respiration lowers photon-use efficiency. Analysis of feasible sub-optimal states identifies a small set of intracellular dissipative motifs. Among them, concomitant operation of the Calvin–Benson cycle and the oxidative pentose phosphate pathway emerges as a parsimonious reaction-level mechanism for dissipative carbon recycling.

Overall, the results provide an experimentally anchored framework linking productive light absorption, energetic coupling, and light respiration to photon-yield losses in a controlled phototrophic system.

## Introduction

Oxygenic phototrophs convert radiant energy into biochemical free energy through photosynthetic electron transport, thereby sustaining carbon fixation and growth. Yet photon-to-biomass conversion efficiencies can vary widely, even within a single organism cultivated under apparently controlled conditions. Explaining these variations at the system level remains difficult because growth emerges from the interplay between the physical structure of the light field, the primary constraints of photochemistry, and the energetic requirements of metabolism.

Two constraints lie at the heart of this problem. First, the photonic resource is not experienced as a uniform external input. In dense phototrophic cultures, radiative attenuation and hydrodynamic mixing expose cells to repeated transitions between strongly contrasted light regimes, so that the relevant light history is spatially heterogeneous and dynamically sampled (Cornet et al., 1998; Cornet and Dussap, 2009; Takache et al., 2010, 2012; Kandilian et al., 2016; Lee et al., 2014). Second, the energetic architecture of oxygenic photosynthesis intrinsically couples ATP production and reducing power generation: proton and electron fluxes remain linked along the photosynthetic electron-transport chain, such that ATP supply per unit electron flow cannot be tuned independently of photochemistry (Alric et al., 2010; Alric, 2014; Kramer and Evans, 2011; Takahashi et al., 2013). Any attempt to interpret growth efficiency must therefore account simultaneously for the physical heterogeneity of the light environment and for the constrained bioenergetic structure of photosynthetic energy conversion.

A key implication is that cells do not respond to incident irradiance *per se*, but to the fraction of absorbed photons that effectively drives reaction-center charge separation and downstream electron transport. Excitations that cannot be processed photochemically are dissipated, notably as heat or fluorescence. Moreover, growth integrates cellular activity over time scales that are far longer than primary photophysical events and typically longer than circulation times in photobioreactors. At the timescale relevant to metabolism and growth, the forcing exerted by light is therefore better described by an effective photochemical input—that is, a time-averaged measure of photochemically productive absorption—than by a boundary irradiance or a local instantaneous light intensity (Cornet and Dussap, 2009; Takache et al., 2012; Kandilian et al., 2016).

Any mechanistic framework for photon-to-biomass efficiency must also address energetic allocation. Net carbon assimilation requires ATP and reducing equivalents in a constrained proportion, while cellular homeostasis, maintenance and redox balancing introduce additional energetic demands. Oxygenic phototrophs therefore rely on regulatory and metabolic degrees of freedom to match photosynthetic supply to intracellular demand (Alric et al., 2010; Alric, 2014; Kramer and Evans, 2011; Takahashi et al., 2013). In this context, light respiration cannot be interpreted a priori as a simple loss term: depending on the energetic regime, it may contribute to balancing ATP and redox budgets, but it may also become dissipative and lower overall photon-use efficiency. Understanding when respiration supports growth and when it penalizes it requires a framework that links reactor-scale light forcing to intracellular energetic organization.

Existing approaches generally illuminate only part of this problem. Radiative-transfer and photobioreactor models describe how incident light is redistributed within dense cultures and how cells experience fluctuating irradiance histories (Cornet et al., 1998; Cornet and Dussap, 2009; Kandilian et al., 2016; Lee et al., 2014), but they usually remain remote from intracellular energetic constraints. Conversely, metabolic and constraint-based approaches describe feasible intracellular flux organizations and energetic trade-offs (Orth et al., 2010; Henry et al., 2007; Beard and Qian, 2005, 2008; Cogne et al., 2011), yet they often represent the light input in a simplified manner and rarely connect it explicitly to a reactor-scale measure of photochemically productive absorption. As a result, the literature still lacks an integrated framework able to relate productive light absorption, energetic partitioning within photosynthetic electron transport, and the role of respiration in shaping photon-to-biomass efficiency.

Here, we propose such a framework for *Chlamydomonas reinhardtii* grown in chemostat photobioreactors. We combine a quantified description of radiative transfer and primary photochemistry with a thermodynamically constrained metabolic analysis in order to connect reactor-scale light forcing to intracellular energetic balances. More specifically, we first define an effective photochemical input, *ϕ*_γ_, based on the rate of photochemically productive photon absorption, and show that the specific growth rate scales linearly with this variable in a near-optimal regime. We then use this experimentally grounded invariant to identify a compatible effective energetic coupling within photosynthetic electron transport and to constrain the admissible balance between photosynthetic activity and respiration. This yields a growth map in the space of photochemical forcing and respiratory activity, allowing a mechanistic interpretation of respiration-limited and dissipative regimes. Finally, we explore the feasible flux space at sub-optimal operation to identify intracellular dissipative motifs compatible with the observed macroscopic signature, including candidate carbon-recycling routes involving the Calvin–Benson cycle and the oxidative pentose phosphate pathway (Farr et al., 1994; Young et al., 2011; Veaudor et al., 2020).

By construction, the present approach does not aim to resolve the full molecular regulation of photosynthetic metabolism. Rather, it seeks an intermediate level of description at which reactor-scale light histories, stoichiometric and thermodynamic constraints, and energetic allocation can be integrated within a common framework. In this way, the study provides an experimentally anchored basis for interpreting photon-yield losses in controlled phototrophic systems and for relating light absorption, bioenergetic coupling and respiration to growth performance.

## Materials and Methods

In this work, *Chlamydomonas reinhardtii* is investigated through a controlled photobioreactor setting that we use as an instrument for quantitative physiology. Accordingly, the Methods combine a quantified description of the culture environment (in particular radiative transfer) with a stoichiometric/energetic modeling framework. The photobioreactor configuration and a major part of the chemostat dataset analyzed here are taken from Takache et al. (2010, 2012). In addition, complementary steady-state data were acquired specifically for the present study on the same experimental setup and using the same protocols. For conciseness, strain description and standard monitoring procedures are not repeated, and the reader is referred to Takache et al. (2010, 2012) for experimental details.

### Radiation field computed by a two-flux formulation

In the present work, the internal light field is computed using a two-flux closure of the radiative transfer equation, which partitions radiation into forward and backward hemispherical components and provides an energetically consistent description of irradiance attenuation in dense, scattering cultures. This approach has been extensively developed and discussed in the photobioreactor literature, primarily for collimated incident light (Cornet et al., 1998; Cornet and Dussap, 2009; Kandilian et al., 2016). Extensions exist to treat partially diffuse or non-collimated illumination (e.g., solar-like conditions) (Lee et al., 2014).

In the particularly common case of a one-sided illuminated planar reactor with a collimated (or quasi-collimated) incident beam, where attenuation can be approximated as onedimensional along the depth coordinate *z*, the two-flux formulation yields an analytically convenient, energetically consistent approximation for the irradiance profile of the form

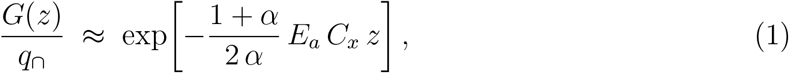

where *q*_∩_ is the incident hemispherical photon flux density (PAR), *C*_*x*_ is the biomass concentration, *E*_*a*_ and *E*_*s*_ are respectively the mass absorption and scattering coefficients, *b* is the backscattered fraction, and

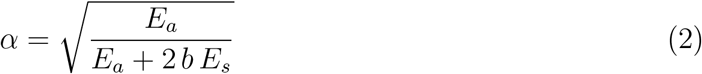

is the corresponding two-flux scattering modulus. Eq. (1) provides a robust reduction of the radiative transfer problem in the operating range relevant to dense, well-mixed photobioreactors (Kandilian et al., 2016).

The profile *G*(*z*) serves as the radiative input for the absorption and primary photochemical conversion steps described next, which define the effective photochemical input used throughout the paper.

### Primary photochemical yield and definition of photochemically productive absorption

To connect the radiative field to metabolic driving, we distinguish (i) photon absorption by the antennae and (ii) the fraction of absorbed excitations that effectively produces reaction-center charge separation. We thus introduce the local biomass-normalized photon absorption (antenna excitation) rate 𝒜 (*z*), obtained from the radiation field as

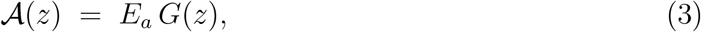

where *G*(*z*) is the local spherical irradiance and *E*_*a*_ the mass absorption coefficient (see Kandilian et al. (2016) for equivalent formulations and conventions).

A classical and physically motivated representation is that the primary photochemical yield decreases with increasing excitation because reaction centers progressively saturate, leading to non-photochemical dissipation and fluorescence. This saturation is commonly captured by hyperbolic laws in photobioreactor modeling (Cornet and Dussap, 2009; Takache et al., 2012). We denote by *ρ*(𝒜) the *primary photochemical yield*, i.e. the fraction of absorbed photons that are photochemically productive at the timescale relevant to growth, and by *ρ*_max_ its maximal value at low excitation:

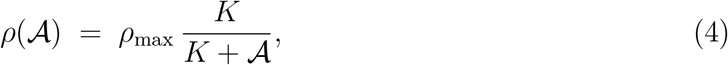

where *K* is the corresponding half-saturation constant on the absorbed-photon scale 𝒜. On this biomass-specific absorption scale, a larger *K* indicates that photochemistry remains close to its low-excitation yield up to higher absorbed-photon loads per unit biomass, i.e. an effectively higher photochemical processing capacity (reaction-center availability and downstream electron-transport/acceptor turnover) before saturation and dissipation dominate. Because 𝒜 is a biomass-specific absorption rate, it provides a cell-centered excitation metric under the physiological conditions considered here (with light as the only possible limiting factor), for which the average cellular mass is expected to vary only weakly (Cogne et al., 2025). In this coarse-grained representation, *K* can be interpreted as an effective photochemical processing-capacity parameter: it aggregates reaction-center turnover together with downstream electron-acceptor availability and competing quenching processes, all expressed on the biomass-specific excitation scale.

The half-saturation constant *K* was taken from the value identified by Takache et al. (2012) for *Chlamydomonas reinhardtii*. In that work, *K* was independently estimated from photosynthetic saturation data using both oxygen-evolution and chlorophyll-fluorescence measurements as functions of irradiance, obtained under standard dilute conditions, i.e. in a regime close to quasi-isoactinic illumination. Because Eq. (4) is written as a function of the biomass-normalized absorption (antenna excitation) rate 𝒜 (Eq. (3)), the same half-saturation level was re-expressed in the units of 𝒜 using the average mass absorption coefficient 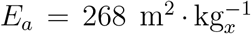 representative of the biomass used in those measurements, yielding:

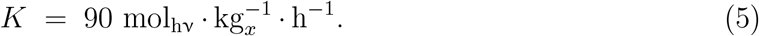

Here, mol_hν_ denotes moles of photons (quanta) in the PAR range.

The maximal primary photochemical yield *ρ*_max_ sets the low-excitation limit of the photochemical conversion law (Eq. (4)) and is often treated as a tunable parameter. In the present work we fix *ρ*_max_ = 0.8 (dimensionless). This value is supported by a second-law upper bound based on radiative exergy under representative solar and aqueous conditions (Appendix A), and is consistent with our LED-based consistency check (Appendix A.3).

Throughout the paper, we denote by γ a *photochemically productive photon*, i.e. an absorbed photon that effectively generates reaction-center charge separation and thus feeds photosynthetic electron transport at the timescale relevant to growth. In contrast, absorbed excitations that cannot be processed by reaction centers are dissipated (heat/fluorescence) and are not counted as γ.

Growth occurs on hour-scale time constants, whereas mixing-induced light fluctuations occur on the order of seconds (typically ∼ 10 s) and primary photophysical events occur on ultrafast time scales (femto-to picoseconds). This separation of time scales supports representing the photochemical forcing relevant to growth by a reactor-scale spatial average of the local rate of photochemically productive absorption. In other words, on the timescale of growth, cells are assumed to sample the full range of local irradiances encountered along the optical path, so that their cumulative photochemical forcing can be approximated by a spatial average.

Finally, we define the non-negative *effective photochemical input ϕ*_γ_ as the reactor-scale average rate of photochemically productive absorption, obtained by combining the local excitation rate 𝒜 (*z*) with the local primary yield *ρ*(𝒜 (*z*)) and averaging over the illuminated depth:

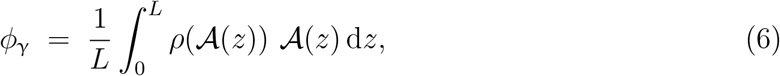

where *L* denotes the illuminated reactor depth.

#### Spectral implementation and reduced formulation

In practice, *ϕ*_γ_ was computed using a spectral implementation over the PAR window, combining the measured source spectrum with wavelength-dependent radiative properties of the culture. For readability, the present manuscript reports the corresponding reduced formulation, written as if *E*_*a*_, *E*_*s*_ and *b* were independent of wavelength, which leads to Eqs. (1)–(6). Implementation details and the associated input data are provided in Supplementary Data S1.

#### Inputs required for optical-property reconstruction

The reconstruction of the wavelength-dependent radiative properties relied on the inverse optical framework of Dauchet et al. (2015) together with the geometric and pigment-related descriptors previously established for *Chlamydomonas reinhardtii* in Takache et al. (2010, 2012). In the range of conditions considered here, the cell-shape model, the cell-size distribution, and the relative pigment distribution were assumed unchanged, consistently with the limited variability reported in those studies. By contrast, the total pigment content was specified for each analyzed steady state and used in the reconstruction of the spectral radiative properties. Biomass concentration was then used separately for the computation of the radiative field within the culture and of the corresponding effective photochemical input *ϕ*_γ_. The corresponding state-specific input data are summarized in Supplementary Data S1.

### Reactor-scale viewpoint and definition of the biotic phase

We view the photobioreactor as an open, multiphase system in which the *abiotic phase* (gas and liquid) surrounds the cells and exchanges matter with the *biotic phase*, defined here as the population of cells present in the reactor. In the following, the biotic phase is assumed homogeneous and represented by average (population-level) quantities, an approximation commonly adopted when the number of cells is large (Schügerl and Bellgardt, 2000).

At the reactor scale, measurable exchange fluxes between the abiotic and biotic phases provide constraints on the net specific transformation rates occurring within the biotic phase; in continuous culture at steady state, this correspondence underlies the link between external exchange rates and intracellular conversion fluxes used throughout the constraint-based analysis.

#### Reactor versus biotic-phase viewpoints for photochemical input

Two equivalent viewpoints are used in the paper.

From a reactor viewpoint, it is convenient to introduce *q*_γ_ as a *reaction rate* representing the net conversion of photochemically productive photons within the biotic phase. In this description, the photochemical input is represented as a biotic-phase conversion rate that acts as a driving term for cellular metabolism.

From a biotic-phase viewpoint, *ϕ*_γ_ ≥ 0 is interpreted as the *effective photochemical input* experienced by the biomass at the timescale relevant to growth. Because cells undergo fast light/dark transitions due to mixing while growth integrates metabolism over much longer timescales, *ϕ*_γ_ is understood as the time-averaged photochemical forcing experienced by cells moving across the heterogeneous light field. In practice, it is computed as the reactor-scale average of photochemically productive absorption (Eq. (6)), where 𝒜 denotes the local antenna excitation (biomass-normalized absorption) rate and *ρ*(𝒜) the corresponding local primary photochemical yield.

Accordingly, under the above sign convention,

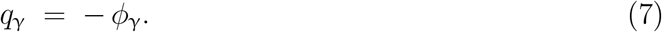

This effective photochemical input is then used as a boundary forcing of the biotic phase in the constraint-based analysis described below.

### Constraint-based metabolic modeling under balanced-growth conditions

We describe the biotic phase as a homogeneous *microreactor* representing the cell population at the reactor scale. Cellular metabolism is represented by a stoichiometric network of *r* intracellular reactions involving *m* compounds. Under the quasi-steady-state approximation (QSSA), intracellular metabolite pools do not accumulate on the timescale relevant to growth. Accordingly, intracellular fluxes satisfy stoichiometric mass-balance constraints (production and consumption rates balance for each intracellular metabolite) (Orth et al., 2010; Nielsen et al., 2003).

Let **J** ∈ ℝ^*r*^ denote the vector of intracellular reaction fluxes (expressed as biomassspecific rates) and let ***ν*** ∈ ℝ^*m*×*r*^ be the stoichiometric matrix. Compounds are partitioned into (i) intracellular metabolites (not exchanged with the medium) and (ii) exchangeable compounds. Accordingly,

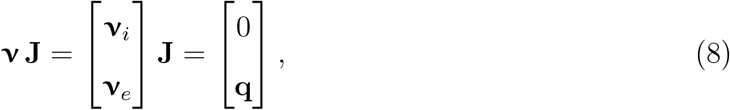

where ***ν***_*i*_ and ***ν***_*e*_ are the submatrices of ***ν*** associated with intracellular and exchangeable compounds, respectively. The vector **q** ∈ ℝ^*p*^ denotes the net exchange rates between the biotic phase and the surrounding abiotic phase, expressed as net conversion rates for exchangeable compounds. Throughout, exchange rates follow the usual algebraic convention: *q*_*k*_ *<* 0 for uptake (consumption from the abiotic phase) and *q*_*k*_ *>* 0 for secretion (production to the abiotic phase).

For phenotype exploration and optimization, it is convenient to treat exchanges as explicit flux variables. Defining the augmented flux vector 𝕁 = [**J**^⊤^, **q**^⊤^]^⊤^, Eq. (8) can be written in the standard extended form

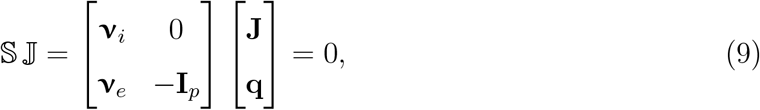

together with bound constraints

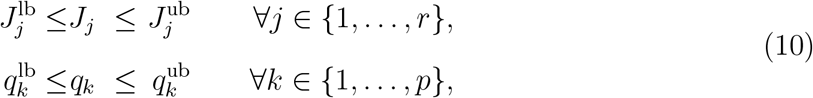

which encode reaction irreversibilities and any additional physiological or experimental information. Here, **I**_*p*_ denotes the *p* × *p* identity matrix, where *p* is the number of exchange fluxes (exchangeable compounds).

### Thermodynamic feasibility constraints

To exclude thermodynamically infeasible flux cycles, we enforce consistency between net flux directions and reaction driving forces, as required by non-equilibrium thermodynamics (Beard and Qian, 2005, 2008; Beard et al., 2002).

Let µ ∈ ℝ^*m*^ denote the vector of chemical potentials of the *m* compounds. Under the ideal-solution approximation, chemical potentials are written as

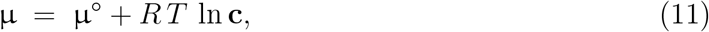

where µ° denotes the vector of reference chemical potentials on the concentration scale (i.e. for *c* = 1 in an ideal solution), **c** is the vector of intracellular concentrations, *R* is the universal gas constant, and *T* is the absolute temperature. The thermodynamic driving forces (affinities) associated with the *r* reactions are then

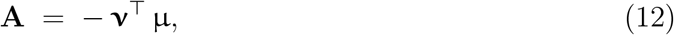

so that *A*_*j*_ is positive when reaction *j* is thermodynamically driven in the forward direction. In addition to reaction-wise sign consistency, thermodynamic feasibility must also hold over internal stoichiometrically closed loops. Let K denote a basis matrix of the null space associated with the internal stoichiometric matrix. The corresponding thermodynamic loop-balance condition is written as

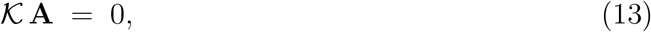

which states that the sum of affinities over any internal stoichiometrically closed loop must vanish, thereby excluding non-physical internal cycles that would generate free energy.

Thermodynamic feasibility further requires that any non-zero net flux proceeds “downhill”, i.e. flux and affinity share the same sign (De Donder-type condition) (Beard and Qian, 2005, 2008):

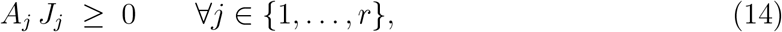

with the understanding that *A*_*j*_ ≠ 0 does not necessarily imply *J*_*j*_ ≠ 0 (e.g. inactive or inhibited enzyme), whereas *A*_*j*_ = 0 enforces *J*_*j*_ = 0.

In practice, Eqs. (13)–(14) are enforced by standard mixed-integer linear constraints that couple the sign of *J*_*j*_ to the sign of *A*_*j*_, using binary variables for forward/reverse operation and a small *ϵ >* 0 to avoid strict inequalities (Henry et al., 2007; Cogne et al., 2011). This ensures that admissible flux solutions do not contain closed internal loops through lumped reactions with no net conversion.

#### Transformed thermodynamic quantities

In the remainder of this work, reaction thermodynamics is evaluated using transformed biochemical quantities defined for equivalent species lumping together all the dissociated forms (Alberty, 1998, 2005), i.e. chemical potentials are understood as 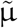 at fixed intracellular *T*, pH and ionic strength. This choice reflects the assumption that intracellular acid–base conditions and ionic strength are effectively regulated (or buffered) on the timescale relevant to growth, so that pH and ionic strength can be treated as prescribed parameters. The corresponding parametrization and standard transformed potentials 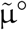 used in the computations are detailed below. For readability, we keep the generic notation µ in Eqs. (11)–(12); numerically, µ is instantiated as 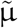 under the above conditions.

### Mathematical formulation

For completeness, the explicit MILP reformulation used to encode the thermodynamic constraints is provided in Appendix B. This reformulation follows the standard linearization strategy introduced in Cogne et al. (2011).

### Thermodynamically constrained flux optimization

To explore admissible steady-state metabolic behaviors, intracellular fluxes are predicted by solving a constraint-based optimization problem following the flux balance analysis (FBA) (Nielsen et al., 2003; Palsson, 2006), subject to the stoichiometric mass-balance constraints, flux and concentration bounds, and the thermodynamic feasibility constraints introduced above (explicit MILP form in Appendix B).

For each imposed set of external operating conditions, fluxes are obtained by solving:

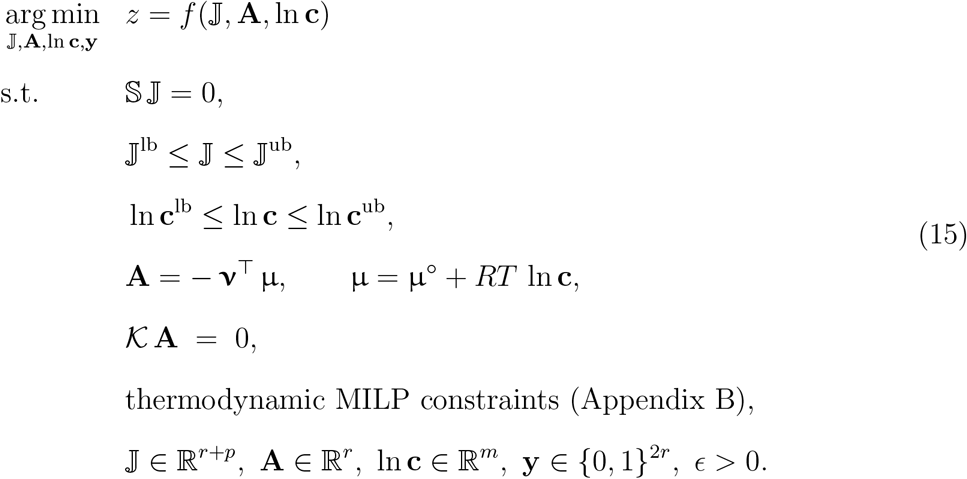

Here, 𝕊 denotes the stoichiometric matrix (including exchange reactions as applicable), 𝕁 the vector of intracellular reaction and exchange fluxes, **c** the vector of intracellular concentrations, and 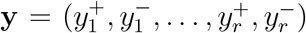 collects the binary direction variables. ***ν*** denotes the intracellular stoichiometric matrix used to compute affinities, **A**. *R* is the universal gas constant and *T* the absolute temperature. The scalar *ϵ* is a fixed tolerance used to replace strict inequalities in the MILP reformulation (Appendix B). The matrix 𝒦 is the null-space basis associated with the internal stoichiometric matrix ***ν***.

Because alternate optima may exist (Mahadevan and Schilling, 2003), we used flux variability analysis (FVA) to determine the admissible range of each reaction flux under optimal operation by minimizing and maximizing each flux while requiring the objective value to remain within a numerical tolerance of its optimum. More generally, the same approach applies under any set of constraints that does not uniquely specify a flux distribution, especially when experimentally motivated operating constraints are incorporated but remain insufficient to fully constrain the system.

#### Computational implementation

All constraint-based optimization problems were implemented in MATLAB R2023b (The MathWorks, Natick, MA, USA). Mixed-integer linear programs arising from the thermodynamic sign-consistency formulation (Appendix B) were solved using the TOMLAB Optimization Environment v8.9 (TOMLAB Optimization Inc.).

### Elementary-process decomposition and *α*-spectra

To interpret feasible flux distributions in terms of pathway-level processes (including exchange/transport), we decomposed steady-state augmented flux vectors 𝕁 = [**J**^⊤^, **q**^⊤^]^⊤^ into conic combinations of a minimal generating set of the flux cone. Under mass-balance together with reaction-direction (irreversibility) and bound constraints, the feasible set of 𝕁 forms a convex polyhedral cone (extended formulation, Eq. (9)). For the metabolic descriptor used here, the imposed reaction-direction constraints (together with bounds) make this cone pointed (i.e. it contains no non-zero lineality space), so it admits a finite minimal generating set given by its extreme rays (Wagner and Urbanczik, 2005). These extreme rays can be viewed as elementary functional processes compatible with network topology, stoichiometry, and reaction directionality (Papin et al., 2002, 2004).

Accordingly, any feasible augmented flux vector 𝕁 can be written as a nonnegative combination of extreme rays {*e*_*ℓ*_}:

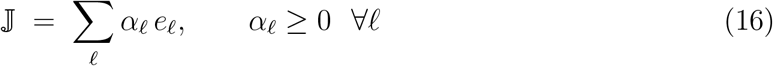

where *α*_*ℓ*_ denotes the contribution (weight) of extreme-ray process *e*_*ℓ*_ to 𝕁.

In general, the decomposition (16) is not unique. For a given feasible state 𝕁, we therefore quantified the admissible range of each weight by computing an *α*-spectrum (Wiback et al., 2003, 2004): for each ray *e*_*ℓ*_, we solved two linear programs that minimize and maximize *α*_*ℓ*_ over ***α*** ≥ 0, subject to (16) and to the same feasibility constraints as the considered flux state (including any additional equalities fixing the operating point). This yields an interval 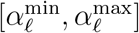 that identifies (i) rays that must contribute 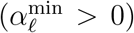 (ii) rays that may contribute 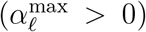 and (iii) rays that cannot contribute 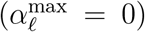. In some conditions, the *α*-spectra collapse to singletons, yielding unique weights for the active set.

Extreme rays were computed using `polco` (Terzer and Stelling, 2008). When needed for cross-checking, elementary modes were computed using `efmtool` (Terzer and Stelling, 2008). *α*-spectra were obtained by linear programming using the same solver stack as for the flux optimization problems.

### Metabolic descriptor and biomass synthesis reaction

#### Network scope and reduction

Cellular metabolism was represented by a stoichiometric network derived from the *Chlamydomonas reinhardtii* model introduced in Cogne et al. (2011). To keep the model computationally tractable while preserving the main degrees of freedom controlling energy/redox partitioning, we used a reduced metabolic descriptor based on network reduction and pathway lumping, consistent with approaches proposed in Rügen et al. (2012). Biosynthetic pathways leading to major biomass building blocks were lumped into a minimal set of lumped biomass-building reactions (proteins, carbohydrates, lipids, pigments, nucleic acids, etc.), whereas central metabolism was retained at the level of elementary reactions. Overall, the resulting metabolic descriptor comprises 86 reactions and 80 metabolites (reactants in the transformed-thermodynamics sense; see below). This reduction preserves the informational content required for quantitative physiology (energy/redox balances and routing flexibility) while limiting the number of explicit metabolites to compounds for which physico-chemical constraints can be consistently defined.

#### Central metabolism representation and compartmentation

The resulting descriptor explicitly includes the main reactions and branch points of central carbon and energy metabolism required to analyze photosynthesis–respiration coupling under autotrophic growth. To maintain a well-posed thermodynamic formulation while keeping the model identifiable from available macroscopic data, compartmentation was represented in a minimal manner. An external compartment was introduced for exchange fluxes with the culture medium, whereas lumped intracellular metabolism was represented by a single effective pool (hereafter referred to as a pseudo-cytosol/chloroplast compartment). Explicit energy-transducing compartments were retained only where electrochemical gradients must be represented, namely the thylakoid lumen for photosynthetic proton translocation and the mitochondrial intermembrane space (IMS) for respiratory proton translocation. This choice avoids introducing poorly constrained transporter fluxes and compartment-specific metabolite pools, while preserving the energy and redox balance requirements needed for the thermodynamically constrained analysis performed here. A schematic representation of the retained network is provided in Fig. 1.

**Figure 1:**
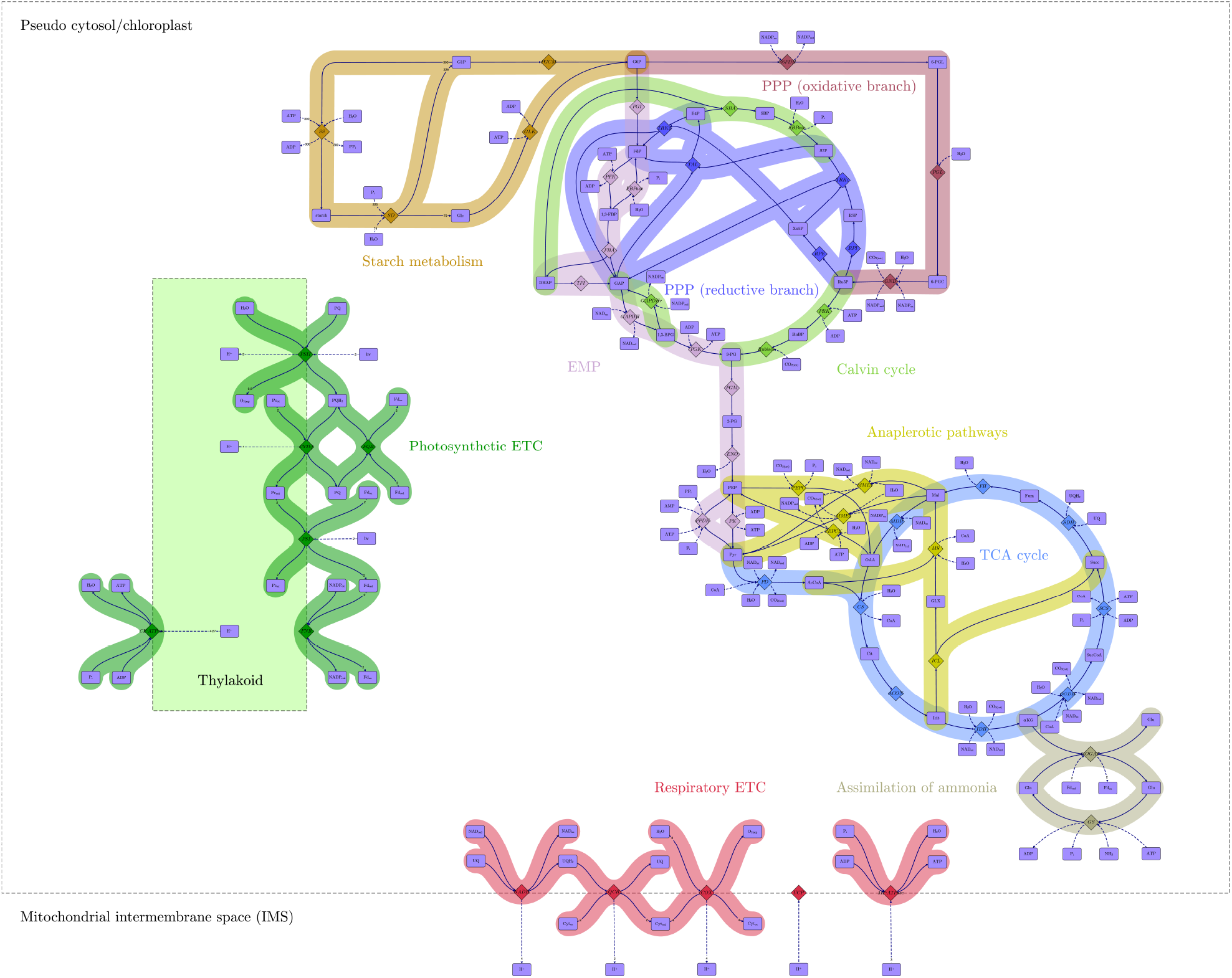
Schematic map of the central metabolism captured by the reduced metabolic descriptor used in this study, including the photosynthetic and respiratory energy-transduction modules used to analyze photosynthesis–respiration coupling under autotrophic growth. Compartmentation is minimal: a single effective intracellular pool (pseudo cytosol/chloroplast) is used for lumped metabolism, while the thylakoid lumen and the mitochondrial intermembrane space (IMS) are retained to represent proton translocation and electrochemical driving forces. Biosynthetic demands are captured in the model via lumped biomass-building reactions but are not detailed in the schematic. The complete descriptor (reaction identifiers, stoichiometry, and metabolite–reaction mapping) is provided as Supplementary Data S2. The map was generated with OMIX (Droste et al., 2011). Abbreviations: TCA, tricarboxylic acid cycle; PPP, pentose phosphate pathway; EMP, Embden–Meyerhof–Parnas pathway (glycolysis); ETC, electron transport chain.

#### Biomass composition and definition of the biomass pseudo-reaction

A biomass synthesis pseudo-reaction was defined from experimentally measured biomass composition representative of *Chlamydomonas reinhardtii* grown under nutrient-replete conditions (Table 1). Macromolecular fractions were determined by standard biochemical assays, while the monomer-level composition used to assemble the biomass pseudo-reaction was taken from Cogne et al. (2011). Measurements were obtained at four chemostat steady states (two incident irradiances and two dilution rates). Since the observed variations remained within measurement uncertainty over the investigated regime, a single average biochemical composition (last column of Table 1) was used to parameterize the biomass reaction. This choice is also consistent with the overall stability of the elemental composition of *Chlamydomonas reinhardtii* biomass produced under varied non-limiting growth conditions, as repeatedly observed over time in our laboratory (Cogne et al., 2025). The resulting pseudo-reaction therefore represents the mean carbon, nitrogen, energy and reductant requirements of balanced growth under nutrient-replete conditions.

**Table 1:**
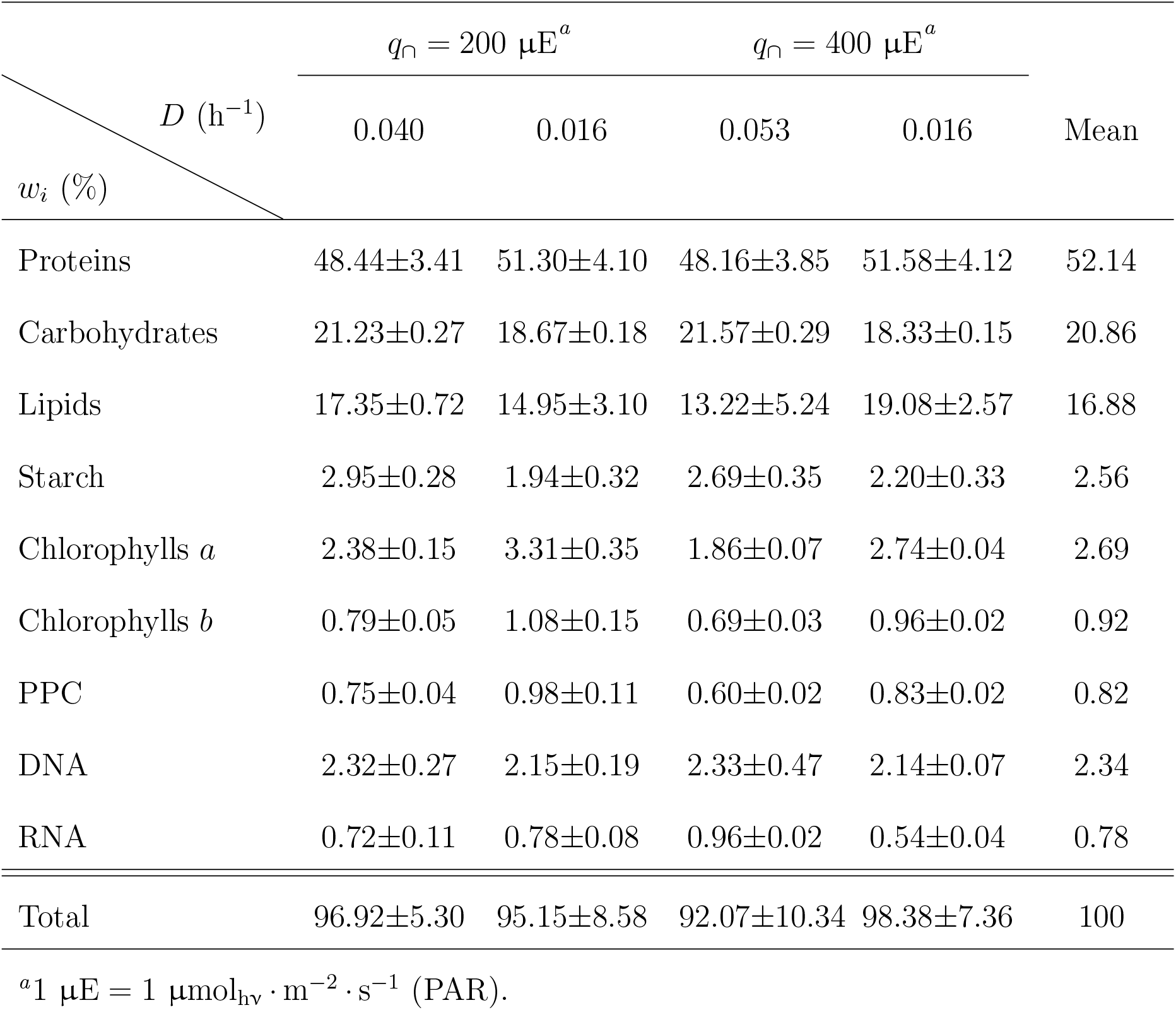
Measured biomass composition at chemostat steady states for two incident photon flux densities (*q*_∩_ = 200 and 400 µmol_hν_ · m^−2^ · s^−1^) and two dilution rates (*D*, in h^−1^). Values are reported as mass fractions *w*_*i*_ of biomass dry weight (% w/w). The last column reports the arithmetic mean over the four conditions; the resulting mean composition was then renormalized to sum to 100%.

#### Normalization, units, and link to chemostat growth rate

For convenience in expressing experimental constraints and interpreting steady-state solutions, the biomass assembly reaction was normalized to the formation of 1 kg of dry biomass. Consequently, all intracellular and exchange fluxes are reported as biomass-specific rates (e.g.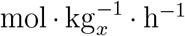). With this normalization, the flux through the biomass reaction, denoted *q*_*x*_, equals the specific growth rate *µ*. Here, the scalar *µ* denotes the growth rate and should not be confused with the vector of chemical potentials µ introduced above.

#### Reactants, species, and transformed potentials

Following biochemical thermodynamics (Beard and Qian, 2005; Alberty, 1998, 2005), we distinguish *species* from *reactants*. A *species* is a chemically distinct form (e.g. a given protonation or metal-complexation state), whereas a *reactant* denotes the biochemical sum of all species of a compound that are rapidly interconverted at fixed *T*, pH and ionic strength. Accordingly, biochemical reactions are written in terms of reactants, while their thermodynamics is captured through transformed equivalent species that lump together all the dissociated forms. In this representation, pH is treated as an independent variable and the contribution of H^+^ is absorbed into transformed standard chemical potentials 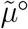 and transformed chemical potentials 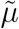 (Beard and Qian, 2005). For instance, the reactant “inorganic carbon” represents the sum of carbonate species 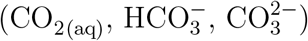 according to the prevailing pH, and its transformed chemical potential corresponds to that biochemical sum.

#### Thermodynamic parametrization and metabolite concentration bounds

All thermodynamic quantities were evaluated using transformed (biochemical) chemical potentials 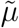 at fixed intracellular conditions *T* = 298 K, pH = 7.5, and ionic strength *I* = 0.25 M (Beard and Qian, 2005; Alberty, 1998, 2005).

In the transformed convention, the bulk-solution proton satisfies 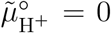. To account for proton translocation across energy-transducing membranes, we introduced compartment-specific transformed standard-state offsets 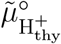 and 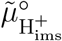 that represent the electrochemical contribution associated with the thylakoid membrane and the mitochondrial inner membrane, respectively. Assuming negligible transmembrane electric potential across the thylakoid membrane (Δ*ψ*_thy_ ≃ 0) and a mitochondrial inner-membrane potential of 150 mV, we set 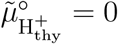 and 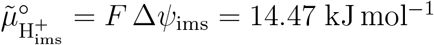, respectively. In problem (15), 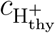 and 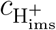 denote effective proton activities encoding the pH differences across these membranes (ΔpH).

The photochemical driving term was represented as an effective standard transformed chemical-potential increment for pigment excitation, 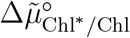, i.e. the free-energy input associated with the transition Chl → Chl^∗^. In practice, this was implemented by introducing an auxiliary photon species γ in the absorption step Chl+γ → Chl^∗^, with 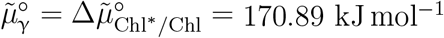 for both photosystems.

Intracellular reactant concentrations were constrained to 10^−10^ ≤ *c*_*i*_ ≤ 10^−2^ M, unless stated otherwise. In addition, the intracellular dissolved oxygen concentration was constrained to exceed the measured O_2_ level in the bulk liquid phase, consistent with net O_2_ production and diffusion out of the biotic phase. Across the experimental conditions analyzed here, the measured dissolved-O_2_ concentration in the bulk liquid ranged from about 150% to 200% of the air-saturation concentration under the operating conditions of the reactor, including aeration, temperature and pressure. At *T* = 298 K and *p* = 1 atm, this corresponds approximately to 0.39–0.52 mM. A single representative mean value of 0.45 mM was therefore retained to define this lower-bound constraint throughout the present analysis.

Reaction identifiers, metabolite identifiers, reaction stoichiometry, and the standard chemical-potential values used in the thermodynamic parametrization are provided in Supplementary Data S2.

#### Bounds on transmembrane pH gradients

The thylakoid pH gradient was constrained to 2 ≤ ΔpH_thy_ ≤ 3.5 (Berg et al., 2002; Ebenhöh et al., 2011), while the pH difference across the mitochondrial inner membrane was constrained to 0 ≤ ΔpH_ims_ ≤ 1 (Nicholls and Ferguson, 2002).

#### Constraint ensuring negligible RuBisCO oxygenation

Photorespiration was assumed negligible under the physiological conditions explored here and was therefore omitted from the model. This assumption is consistent with the criterion of Kazbar et al. (2019), based on the dissolved inorganic carbon-to-oxygen ratio in the reactor, which was monitored and controlled through gas-phase operating conditions (Takache et al., 2010). To enforce this assumption in the thermodynamically constrained formulation, we imposed an additional constraint on the local O_2_*/*CO_2_ ratio governing the competition between the carboxylase and oxygenase activities of RuBisCO:

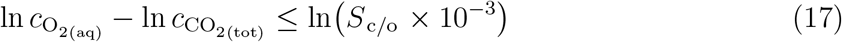

where 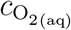 denotes the concentration of dissolved oxygen and 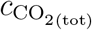 denotes the total inorganic carbon *reactant* available to RuBisCO, i.e. the biochemical sum of dissolved inorganic carbon species under the prevailing intracellular conditions, and *S*_c/o_ = 61 is the relative selectivity factor of *Chlamydomonas* RuBisCO for CO_2_ over O_2_ (Badger et al., 1998). Equation (17) constrains RuBisCO oxygenase activity to remain below 0.1% of carboxylase activity, i.e. within a range treated here as negligible.

## Results

### An experimentally grounded invariant links growth to photochemically productive light

We first established an experimentally grounded relationship linking growth to the fraction of incident light that is effectively absorbed and converted into photochemistry at the timescale of growth (hereafter referred to as the *effective photochemical input*). We considered a reference regime of near-optimal operation in continuous culture, defined operationally as the set of chemostat steady states that maximize volumetric biomass productivity at each imposed incident photon flux density *q*_∩_ for a rectangular photobioreactor illuminated from one side by a collimated light source (Takache et al., 2010). In practice, for each *q*_∩_ this regime is characterized by an optimal steady-state growth rate *µ*_opt_ (equal to the optimal dilution rate *D*_opt_).

For each *q*_∩_, we extracted *D*_opt_ and computed the associated effective photochemical input *ϕ*_γ_ to the biotic phase (Table 2). While *D*_opt_ follows directly from the optimal chemostat steady state, *ϕ*_γ_ is a derived quantity. For each steady state, the internal light field was characterized by a spectral light-transport calculation parameterized with biomass optical properties (absorption/scattering) reconstructed for that condition using the inverse optical-property method of Dauchet et al. (2015). *ϕ*_γ_ was then obtained by combining this field with the local primary photochemical yield *ρ*(𝒜) (reaction-center saturation) and integrating over the culture volume (Eqs. (1)–(6)), as defined in Materials and Methods.

**Table 2:** Optimal dilution rates (*D*_opt_) and effective photochemical inputs (*ϕ*_γ_) determined from steady-state *C. reinhardtii* chemostat cultures for different imposed incident photon flux densities (*q*_∩_) (Takache et al., 2012).

| $q_{\square}$ ($\mu\text{mol}_{\text{hv}} \cdot \text{m}^{-2} \cdot \text{s}^{-1}$) | $D_{\text{opt}}$ ($\text{h}^{-1}$) | $\phi_{\gamma}^a$ ($\text{mol}_{\gamma} \cdot \text{kg}_x^{-1} \cdot \text{h}^{-1}$) |
| --- | --- | --- |
| 110 | 0.032 | 14.16 |
| 200 | 0.041 | 19.15 |
| 300 | 0.054 | 22.51 |
| 400 | 0.057 | 25.26 |
| 500 | 0.058 | 26.59 |
| 600 | 0.061 | 28.38 |

<sup>a</sup> Re-expressed from Takache et al. (2012).

Across the reference regime, the optimal growth rate scales approximately linearly with *ϕ*_γ_ (Fig. 2), indicating an operationally constant biomass yield on photochemically productive photons. A regression of *µ*_opt_ against *ϕ*_γ_ using a linear model without intercept yields a high goodness of fit (*R*^2^ = 0.96) and the estimate

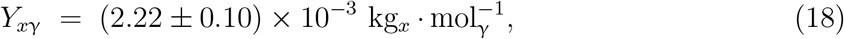

so that, within this reference regime,

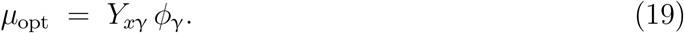

**Figure 2:**
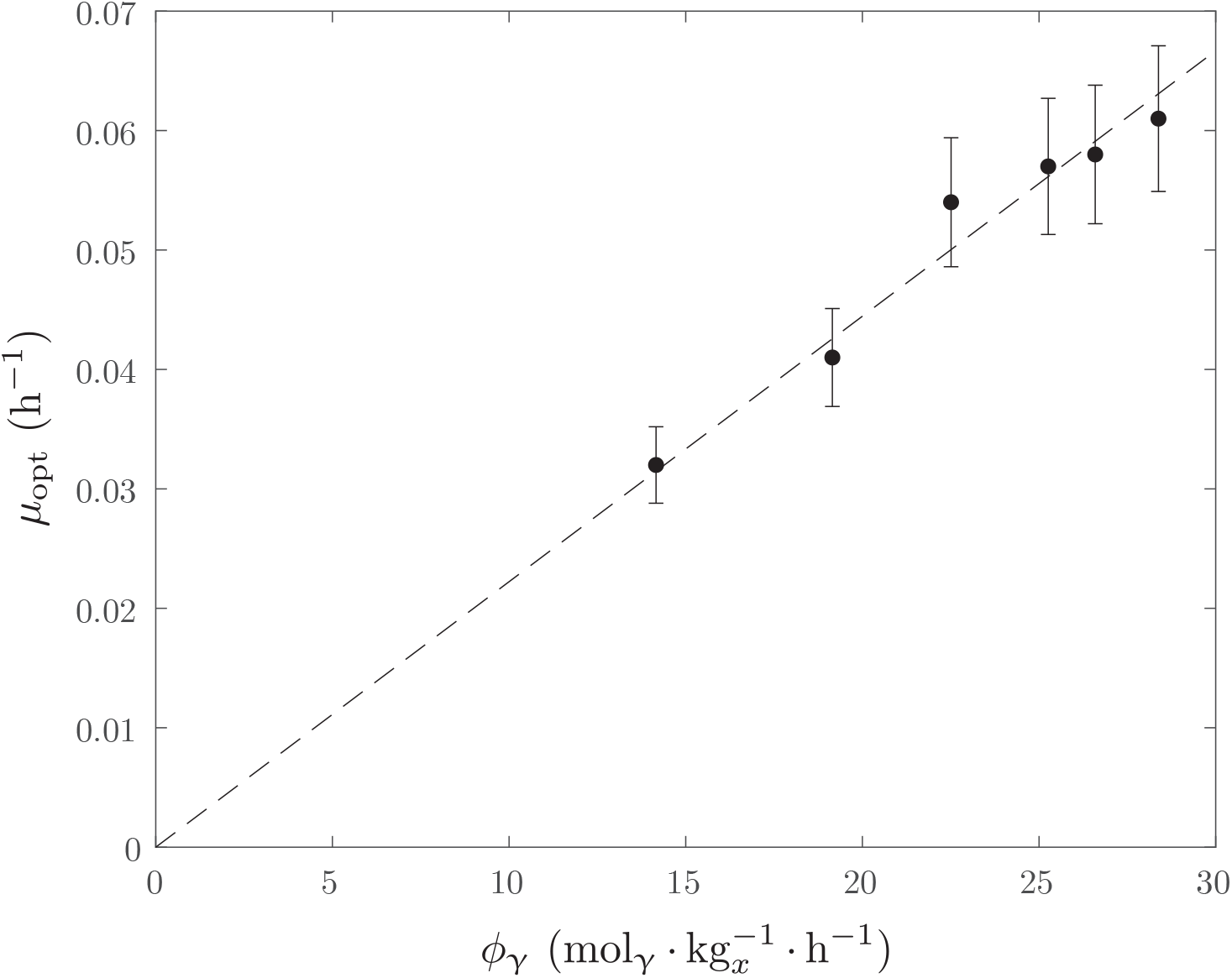
Relationship between the optimal steady-state growth rate and the effective photochemical input in the near-optimal chemostat regime. Each point corresponds to a chemostat steady state maximizing volumetric biomass productivity at a given imposed incident photon flux density *q*_∩_ (Table 2) (Takache et al., 2010). The effective photochemical input *ϕ*_γ_ was estimated for each operating point from the light-transport and primary photochemical conversion model (Eqs. (1)–(6)). The solid line shows a linear regression constrained to pass through the origin, yielding *µ*_opt_ = *Y*_*x*γ_ *ϕ*_γ_ with 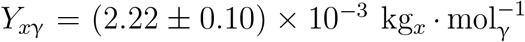. Error bars correspond to an assumed relative uncertainty of 10% on *D*_opt_, introduced here for illustrative sensitivity purposes in the absence of explicit experimental uncertainty estimates in Takache et al. (2012).

The absence of a statistically supported intercept is consistent, over the explored operating range and under this operational definition of near-optimal operation, with a negligible apparent non-growth-associated energetic demand when growth is expressed against *ϕ*_γ_. In the next section, we impose Eq. (19) to calibrate the effective photosynthetic energetic coupling such that the model reproduces the steady-state growth–photochemical-input relationship observed in chemostat (Takache et al., 2010, 2012).

### Inferring an effective photosynthetic energetic coupling under growthoptimal operation

We next used the reference constraint established above to infer an energetically efficient coupling within photosynthetic electron transport, summarized by an effective ratio ATP / 2e^−^ (ATP synthesized per two electrons transferred through photosynthetic electron transport). The underlying idea is that, for a fixed stoichiometric description of cellular metabolism operated at steady state, the partitioning of photochemical energy between ATP supply and reductant generation determines whether the experimentally observed scaling between growth and effective photochemical input can be met without invoking additional, purely dissipative fluxes.

Throughout this analysis, the respiratory module was evaluated under an efficiently coupled baseline. Because O_2_ is in excess due to net photosynthetic production, terminal electron acceptor limitation was neglected in the studied regime; we therefore constrained oxidative phosphorylation to operate near maximal coupling and fixed the effective phosphorylation yields to P/O = 2.5 for NADH-supported entry and P/O = 1.5 for FADH_2_/succinatesupported entry (Hinkle, 2005; Divakaruni and Brand, 2011; see Appendix C for the corresponding linear constraint). Any additional uncoupling, if required by feasibility in specific scenarios, is represented explicitly through the proton-leak degrees of freedom already included in the descriptor.

Specifically, we imposed the experimentally grounded relationship linking growth to effective photochemical input (Eq. (19)) in the signed-flux formalism as

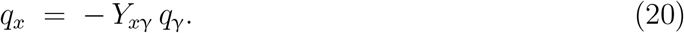

We then sought an effective photosynthetic energetic coupling, denoted *κ* ≡ ATP / 2e^−^, such that steady-state solutions remain feasible under Eq. (20) without requiring the generic ATP-hydrolysis dissipation sink.

More precisely, for a given candidate coupling *κ*, we computed the maximum admissible flux through the ATP-hydrolysis sink, 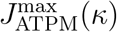, i.e. the largest purely dissipative ATP demand that can be accommodated while still satisfying Eq. (20) under the model constraints: 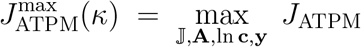 s.t. model constraints, Eq. (20), and coupling *κ* fixed.

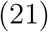

The quantity 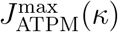 can be interpreted as a dissipative slack, i.e. the largest purely dissipative ATP-hydrolysis flux compatible with Eq. (20) under the model constraints. In our parameterization, increasing *κ* corresponds to increasing the contribution of cyclic electron flow, which raises ATP supply per PSII-supplied (linear) electron while reducing the net availability of reductant for biosynthesis. As a result, the admissible ATP-dissipation slack 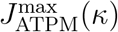 decreases with *κ* and reaches zero at a threshold *κ*^⋆^.

We therefore define the threshold coupling *κ*^⋆^ as the largest value of *κ* for which Eq. (20) remains feasible, and for which feasibility requires an inactive ATP dissipation sink, i.e. 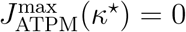, meaning that no feasible solution exists with *J*_ATPM_ *>* 0 (and thus *J*_ATPM_ = 0 in all feasible solutions). For *κ > κ*^⋆^, Eq. (20) becomes infeasible under the model constraints, indicating that the imposed growth–photochemical-input scaling cannot be met because reductant supply becomes limiting.

Equivalently, *κ*^⋆^ can be obtained by maximizing the photosynthetic ATP yield per unit PSII electron supply under the same constraints and under Eq. (20), which leads to the following linear-fractional program:

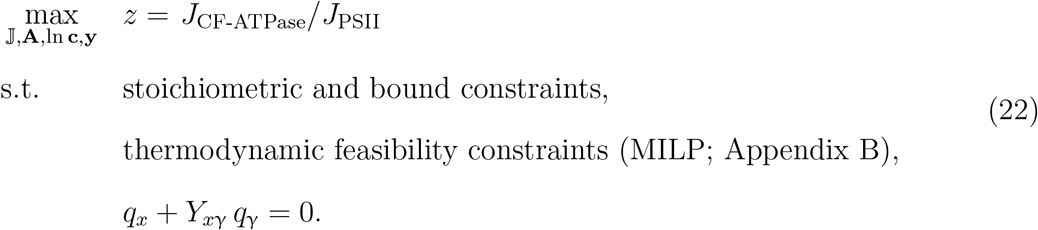

Here, *J*_CF-ATPase_ and *J*_PSII_ denote the fluxes through the photosynthetic ATP synthase and the PSII water-oxidation reaction (electron supply), respectively. The linear-fractional objective was handled using the standard reformulation strategy detailed in Cogne et al. (2011). This yields an effective coupling ATP / 2e^−^ = 1.45 ± 0.15, which we interpret as a cell-scale coupling consistent with near-optimal growth in the considered regime.

A key outcome is that, at this coupling, the reference regime can be reproduced without requiring a non-zero flux through the generic ATP-hydrolysis sink introduced as an explicit dissipative demand (“maintenance”). In other words, the inferred coupling reconciles the observed proportionality between growth and effective photochemical input using only biosynthesis and the redox/energy balancing already captured by the stoichiometric network, without introducing an *ad hoc* energetic drain. This result is consistent with the energetic flexibility of green algae, where cyclic electron flow and related processes provide a lever to tune ATP supply relative to reductant production (Alric et al., 2010; Alric, 2014).

In the remainder of the study, we fix ATP / 2e^−^ to this inferred value, i.e. the largest coupling compatible with the reference growth–photochemical-input scaling while keeping the ATP-hydrolysis sink inactive (respiratory P/O fixed as above).

### Two reconstructed chemostat states reveal a loss of identifiability under sub-optimal yield

Using the constraint-based thermodynamic framework described in Materials and Methods, we analyzed two chemostat steady states at *q*_∩_ = 200 µmol_hν_ · m^−2^ · s^−1^. Throughout, the effective photosynthetic energetic coupling was fixed to the inferred value ATP / 2e^−^ = 1.45, and the respiratory module was kept at the same efficiently coupled baseline with fixed P/O ratios (linear energetic-coupling constraints detailed in Appendix C). This common energetic parameterization is consistent with the near-constant biomass composition observed under nutrient-replete balanced-growth conditions and with the fact that oxygen remains in excess for respiration in all analyzed states.

For each state, we imposed (i) the measured biomass synthesis flux *q*_*x*_ = *µ* = *D* and (ii) the corresponding photochemical forcing *q*_γ_ (derived from the optical and primary photochemical calculations described in Materials and Methods). We then characterized the admissible intracellular flux space by flux variability analysis (FVA): each reaction flux *J*_*j*_ was successively minimized and maximized under the same set of stoichiometric, thermodynamic and bound constraints, while maintaining the imposed (*q*_*x*_, *q*_γ_) operating point.

The first state corresponds to a new chemostat experiment specifically designed to reproduce, at *q*_∩_ = 200 µmol_hν_ · m^−2^ · s^−1^, the operationally optimal regime previously identified from the Takache criterion (Takache et al., 2010). In this experiment, biomass productivity reached a maximum at *D* = 0.041 h^−1^, with the corresponding photochemical forcing magnitude 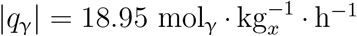 (Martzolff, 2013). Under the fixed energetic parameterization retained here, this reconstructed state is consistent with the solution expected to maximize the objective function *z* = −*q*_*x*_*/q*_γ_ under the imposed constraints, i.e. the biomass yield on photochemically productive photons *Y*_*x*γ_, consistently with Cogne et al. (2011). FVA returned extremely narrow admissible ranges around this operating point, indicating that the reconstructed near-optimal state is essentially unique under the imposed constraints. Moreover, solving the optimization problem in the spirit of Cogne et al. (2011), under the experimentally derived upper bound on biomass synthesis flux and with *z* = −*q*_*x*_*/q*_γ_ as objective, yielded 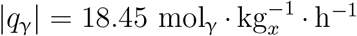, i.e. within less than 3% of the value estimated experimentally. This close agreement, together with the near-uniqueness revealed by FVA, justified retaining the corresponding optimal solution as the reference state in the subsequent pathway-level analysis; this reference state is shown in Fig. 3. It is also consistent with the metabolic organization previously inferred for growth-optimal operation in Cogne et al. (2011), in particular an open TCA cycle and the absence of detectable oxidative pentose phosphate pathway (oxPPP) recruitment. In this regime, cultures operate in the “fully light-absorbing” configuration described by Takache et al. (2012), where essentially all incident PAR is absorbed across the reactor depth while avoiding a persistent dark zone, i.e. without a permanently light-deprived region at reactor scale.

**Figure 3:**
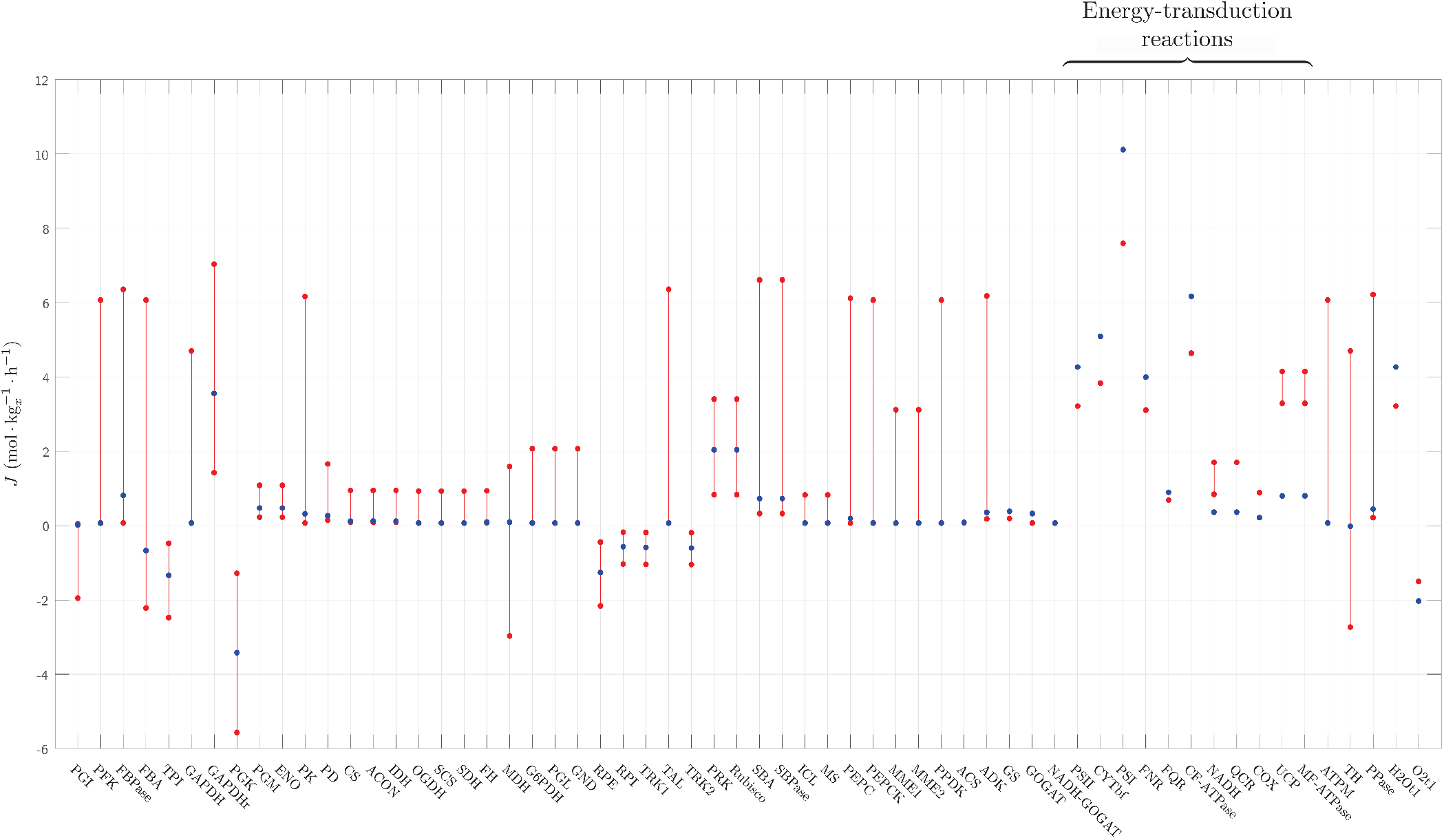
Flux variability analysis (FVA) for two chemostat steady states at *q*_∩_ = 200 µmol_hν_ · m^−2^ · s^−1^. For each state, the measured growth rate (*q*_*x*_ = *µ* = *D*) and corresponding photochemical forcing (*q*_γ_) were imposed, with ATP / 2e^−^ = 1.45 and an efficiently coupled respiratory baseline (fixed P/O ratios). Bars report the admissible ranges of each intracellular flux *J*_*j*_ obtained by successively minimizing and maximizing *J*_*j*_ under the same stoichiometric, thermodynamic and bound constraints. Under the near-optimal reference state (*D* = 0.041 h^−1^), the solution is essentially unique (narrow ranges), whereas under the sub-optimal condition (*D* = 0.016 h^−1^; stronger self-shading), many central-metabolism fluxes become poorly identified while photosynthetic and respiratory energy-transduction fluxes remain tightly constrained, notably *J*_PSII_ and *J*_COX_. Reaction identifiers and corresponding biochemical assignments are provided in Supplementary Data S2.

The second state was obtained under the same incident irradiance and experimental setup, but at a lower dilution rate *D* = 0.016 h^−1^, yielding a photochemical forcing magnitude 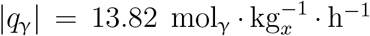 (Martzolff, 2013). This condition leads to a higher biomass concentration (stronger self-shading) and to an apparent yield on photochemically productive photons, 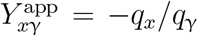, that is lower than the reference invariant *Y*_*x*γ_ established under near-optimal operation. Under this sub-optimal condition, the inverse problem becomes underdetermined: many central-metabolism fluxes span broad admissible ranges (Fig. 3).

Importantly, the two reconstructed states differ markedly in identifiability: the near-optimal state is essentially unique under the imposed constraints, whereas the sub-optimal state remains underdetermined in central metabolism. Despite this loss of identifiability, the photosynthetic and respiratory energy-transduction fluxes remain tightly constrained across the admissible set. This is a direct consequence of fixing the effective photosynthetic energetic coupling ATP / 2e^−^ together with the respiratory P/O ratios: once the energetic partitioning within photosynthetic electron transport and the respiratory ATP return are prescribed, the energy- and redox-balance constraints embedded in the stoichiometric network strongly restrict the admissible photosynthesis–respiration balance. As a result, both the terminal oxidase flux *J*_COX_ and the PSII water-oxidation flux *J*_PSII_ are well identified even in the sub-optimal condition, enabling a robust positioning of both states in the photosynthesis–respiration space.

Notably, moving from the reconstructed near-optimal to the sub-optimal condition, the ratio *J*_COX_*/J*_PSII_ increases, indicating a higher respiratory demand per unit photosynthetic electron supply under stronger self-shading.

### A two-dimensional growth map reveals respiration-limited and dissipative regimes

With the effective photosynthetic coupling fixed, the photochemical conversion rate *q*_γ_ becomes a direct control parameter for photosynthetic energy supply to metabolism: increasing photochemically productive light scales the delivery of ATP and reducing power through photosynthetic electron transport. We then examined how growth is shaped by the joint variation of photosynthetic supply and respiratory activity.

Respiratory activity was parameterized by the terminal cytochrome oxidase flux *J*_COX_, used here as a proxy for mitochondrial respiratory electron flow. Scanning a physiologically relevant range of pairs (*q*_γ_, *J*_COX_), we computed the *growth potential* supported by each pair under the stoichiometric, energetic and thermodynamic constraints of the model. This yields a two-dimensional map 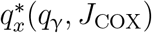 over the photosynthesis–respiration space (Fig. 4), where 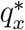 denotes the highest admissible biomass production rate compatible with the constraints for the considered (*q*_γ_, *J*_COX_). Under the biomass-specific normalization used here, *q*_*x*_ coincides with the specific growth rate *µ*.

**Figure 4:**
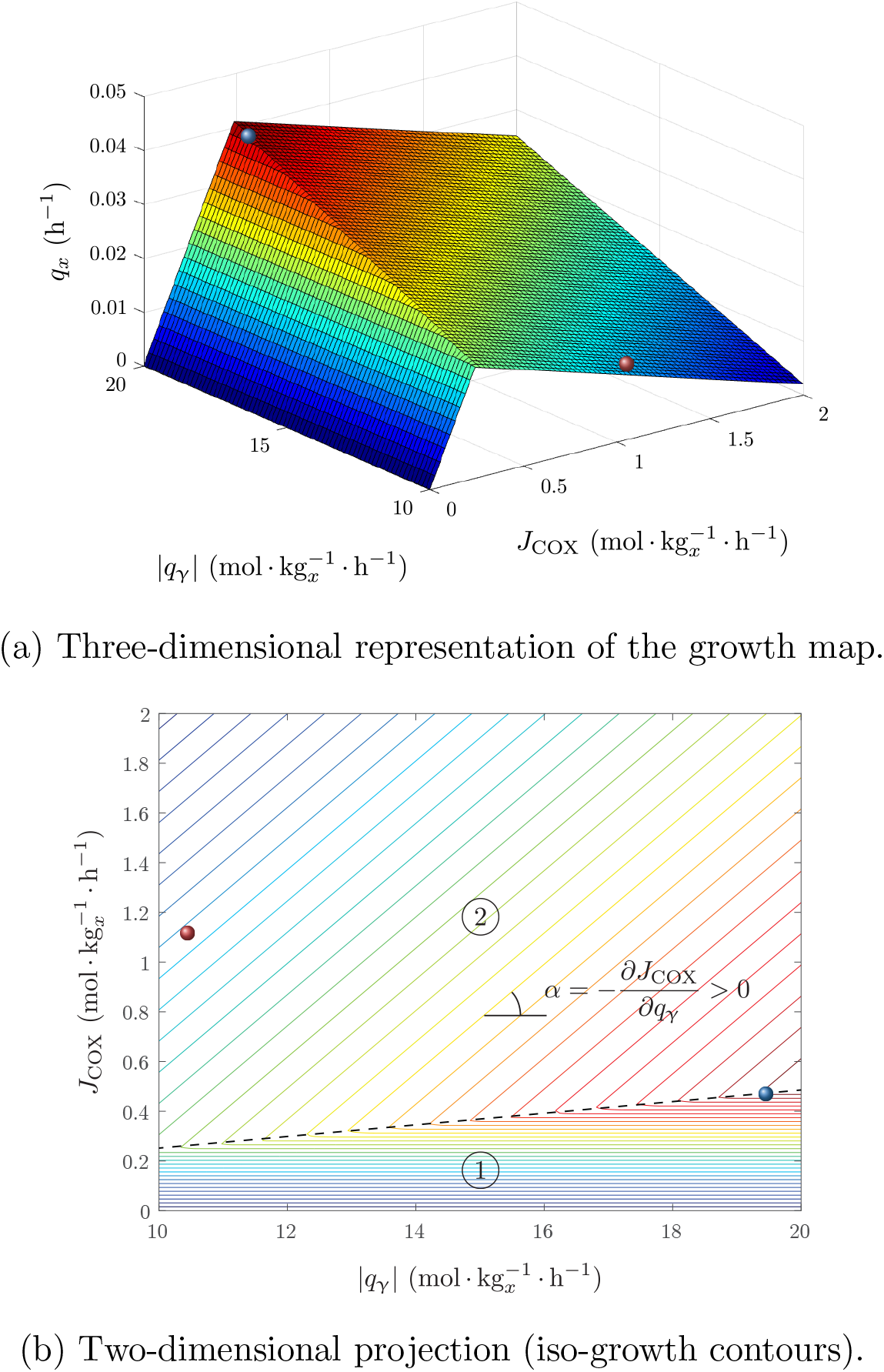
Two-dimensional growth map in the photosynthesis–respiration space. The surface (a) and contour projection (b) show the growth potential 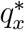 as a function of the photochemical conversion rate *q*_γ_ and the terminal oxidase flux *J*_COX_, with ATP / 2e^−^ fixed to the inferred value and respiration defined by an efficiently coupled baseline (fixed P/O ratios). Region I corresponds to a respiration-limited regime, whereas Region II defines a dissipative (over-respiring) regime in which higher respiratory flux requires higher photochemical input to sustain the same growth. The dashed line denotes the locus of maximal growth potential separating both regions. The operationally optimal reference state 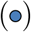 and the sub-optimal state under stronger self-shading 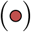 are shown for comparison.

The two chemostat states analyzed above are also projected onto this map to relate experimentally observed operation to the predicted regime structure: the operationally optimal reference state at 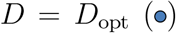 and the sub-optimal, stronger self-shading state at 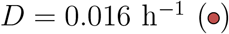.

The map reveals two qualitatively distinct regions. In Region I, iso-growth contours are approximately horizontal in the (*q*_γ_, *J*_COX_) plane: increasing respiratory flux increases the growth potential at essentially fixed photochemical input, indicating a *respiration-limited regime*. In Region II, iso-growth contours display a positive slope: maintaining a given growth rate requires increasing photochemical input as respiratory flux increases. We therefore interpret Region II operationally as a *dissipative (over-respiring) regime*, because additional respiratory activity incurs an increased photochemical demand for the same macroscopic outcome.

A ridge of maximal growth potential delineates the set of (*q*_γ_, *J*_COX_) combinations that maximize 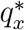 under the imposed constraints. Along this ridge, the photosynthesis–respiration partition is well approximated by a nearly constant respiratory demand relative to photosynthetic oxygen evolution. On an O_2_ molar basis, we obtain

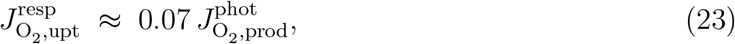

where 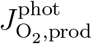 and 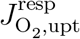 denote, respectively, the photosynthetic oxygen production flux and the respiratory oxygen uptake flux. Departures from this balance, either toward lower or higher respiratory activity at fixed photochemical input, result in a marked decrease in growth potential.

Taken together, these results suggest that the maximal-growth ridge defines a one-parameter family of growth-optimal steady states in the (*q*_γ_, *J*_COX_) plane: for each photochemical conversion rate *q*_γ_, there exists an optimal respiration level that maximizes 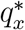 under stoichiometric, thermodynamic, and bioenergetic constraints, including limits imposed by ATP yield per electron transferred (P/2e^−^ in photosynthesis and P/O ratios in respiration).

In this view, the maximal-growth ridge can be interpreted as the locus of growth-optimal operating points that, for a given *q*_∩_, connects a *kinetic regime*—where photochemical input is in excess and a non-negligible fraction of the incident PAR is transmitted through the culture—to the onset of a *radiative-transport-limited regime*, where incident PAR is essentially fully absorbed across the illuminated depth and effectively redistributed over an increasing biomass concentration (Cornet et al., 1998; Cornet and Dussap, 2009; Takache et al., 2010). Beyond this transition, further increases in biomass concentration primarily strengthen self-shading and lower the effective photochemical input available to growth.

Accordingly, for fixed *q*_∩_, decreasing *D* increases self-shading and reduces the effective photochemical conversion rate, thereby moving the system away from the maximal-growth ridge.

Finally, the operationally sub-optimal steady state 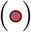 falls within the dissipative region, whereas the operationally optimal reference state 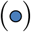 lies close to the maximal-growth ridge. This supports the interpretation that the ridge captures, in the (*q*_γ_, *J*_COX_) space, the family of steady states compatible with the near-optimal chemostat regime defined operationally by Takache et al. (2012).

### Identifying dissipative pathway motifs compatible with dissipative states

To move beyond regime classification, we sought intracellular mechanisms capable of realizing the macroscopic signature of dissipative operation highlighted by Fig. 4, namely an increased photochemical demand at unchanged growth when the photosynthesis–respiration balance shifts toward higher respiratory activity. We therefore analyzed the operationally sub-optimal chemostat steady state introduced above, which lies within the dissipative region.

All bioenergetic parameters were kept fixed to the values inferred previously (photosynthetic coupling *κ*^⋆^ = ATP / 2e^−^ and an efficiently coupled respiratory baseline), and the operating point was specified by the experimental constraints *q*_*x*_ = *D* = 0.016 h^−1^ and 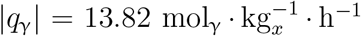. Under these constraints, the admissible steady-state solution set remains non-unique for large parts of central metabolism (Fig. 3), motivating a systematic search for representative dissipative flux patterns. Under the imposed equality and bound constraints, the corresponding feasible steady-state set forms a bounded polytope in flux space, so that its vertices can be interpreted as extreme admissible metabolic states.

We interrogated this feasible set by enumerating its vertices (extreme feasible states). Vertices capture extreme realizations of the active constraints and often reveal qualitatively distinct operating regimes, i.e. flux patterns that cannot be expressed as convex combinations of other feasible states. Each vertex was therefore treated as an extreme admissible metabolic state compatible with the considered operating point. This yielded 224 vertices. We then filtered the vertex set for thermodynamic consistency by enforcing the thermodynamic feasibility constraints described in Materials and Methods and discarding vertices that imply thermodynamically infeasible internal cycles, as in Rügen et al. (2012). After this filtering, 112 thermodynamically admissible vertices remained, corresponding to 112 distinct thermodynamically feasible flux states under the considered operating point.

Finally, the remaining thermodynamically feasible vertex states were interpreted at the pathway level by decomposition into elementary processes. Specifically, we considered the associated steady-state flux cone defined by the same stoichiometric, directionality, bioenergetic and bound constraints, but without the exchange-flux equalities used to close the polytope at the chemostat operating point. Each filtered vertex flux distribution was then expressed as a nonnegative combination of extreme rays of this cone (Methods), enabling identification of underlying elementary processes and recurring dissipative motifs.

In practice, the reduced descriptor admits 261 elementary modes. Because the corresponding steady-state flux cone is pointed, its non-redundant generating set is given by its extreme rays. We computed 42 such rays, which define 42 elementary processes that are necessary and sufficient to generate all thermodynamically feasible flux states of the model under the imposed directionality constraints. For clarity, these processes are indexed arbitrarily from 1 to 42, and the ordering carries no meaning. Where relevant, potential non-uniqueness in the decomposition of a given flux state was quantified by *α*-spectra (Methods).

In practice, the decomposition was unique for almost all flux states analyzed here, including the growth-optimal reference state. This greatly facilitates pathway-level interpretation. Although no formal proof is provided, this near-systematic uniqueness likely reflects the strong restriction of energetic degrees of freedom induced by the imposed photosynthetic and respiratory coupling constraints, in particular the fixed P / 2e^−^ and P/O ratios.

To further reduce the space of candidate dissipative mechanisms, we restricted attention to extreme feasible states whose elementary-process decomposition contains the same biomass-producing process, denoted *e*_11_. Operationally, *e*_11_ was identified as the unique extreme ray with non-zero weight in the decomposition of the growth-optimal reference state that carries the biomass synthesis reaction. It was therefore used here as a *growth core* to compare optimal and sub-optimal operation. The overall workflow is summarized in Fig. 5.

**Figure 5:**
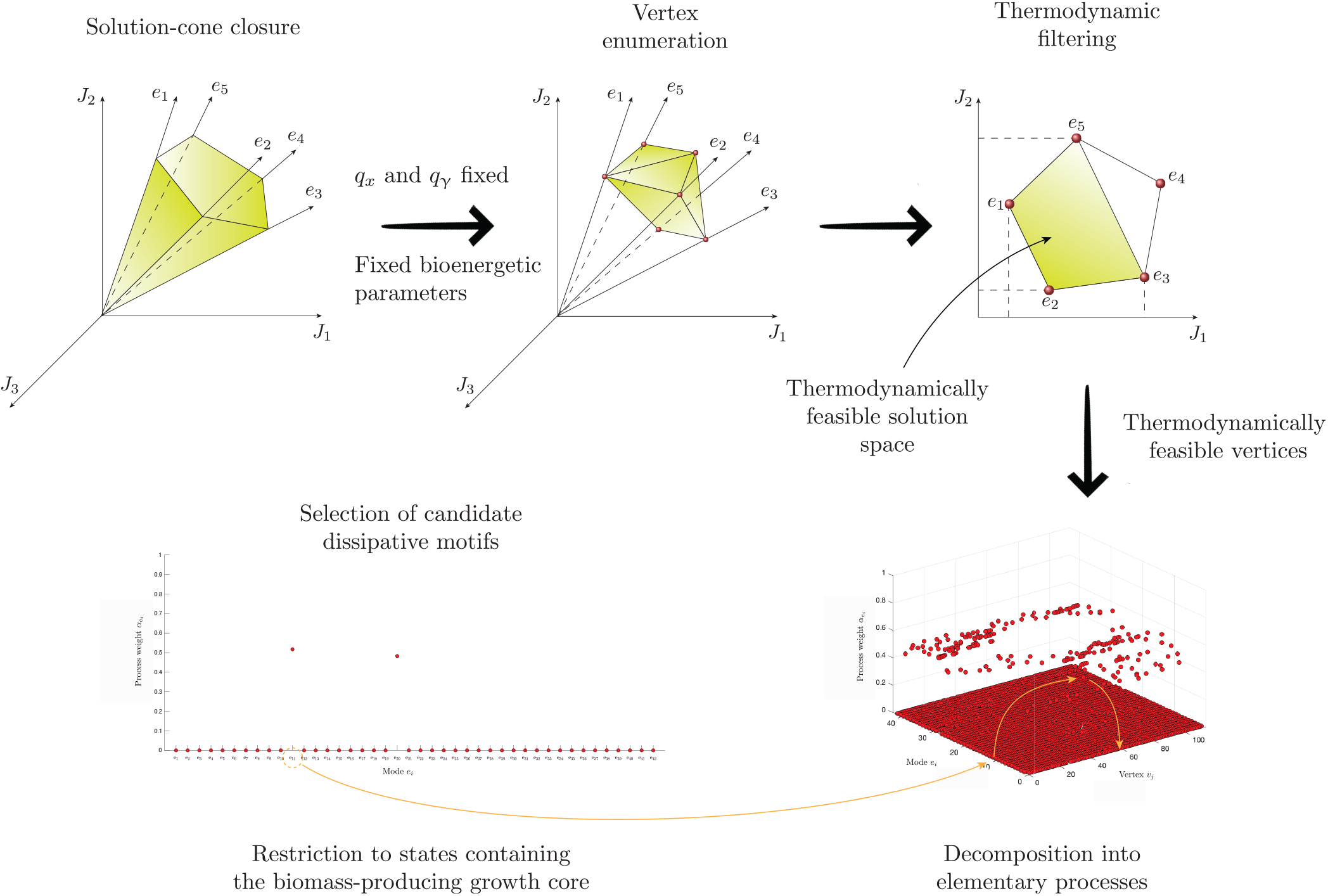
Workflow for identifying dissipative pathway motifs compatible with the operationally sub-optimal chemostat state. From the experimentally constrained dissipative operating point (*q*_*x*_ = *D* = 0.016 h^−1^ and 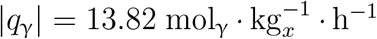), the feasible polytope was obtained by closing the corresponding steady-state flux cone under fixed bioenergetic parameters. Extreme feasible states were then enumerated, filtered for thermodynamic consistency, and decomposed into extreme rays of the associated flux cone to identify the underlying elementary processes. Candidate dissipative motifs were finally selected by retaining only states containing the biomass-producing growth core *e*_11_.

Six extreme feasible states representative of sub-optimal (dissipative) operation were identified and characterized under the same operating-point constraints. Each of these states admits a sparse extreme-ray decomposition involving only two elementary processes: the biomass-carrying process *e*_11_ and a second, state-specific process associated with the futile behavior described above. This defines a common growth backbone (*e*_11_) onto which distinct dissipative motifs are superimposed (Fig. 6).

**Figure 6:**
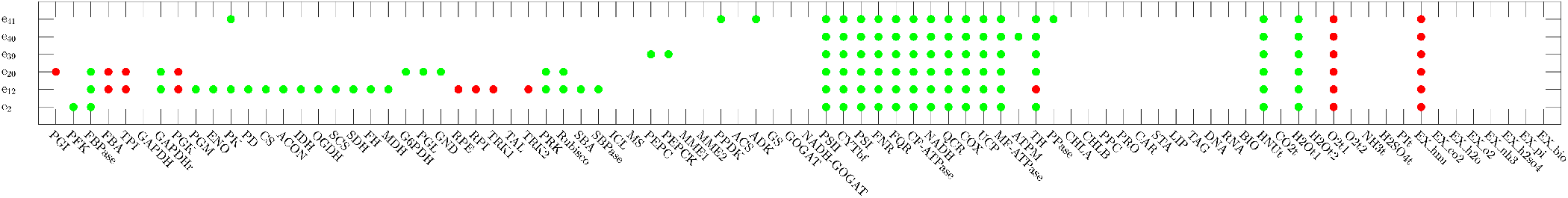
Reaction-level representation of the six candidate dissipative processes identified from the decomposition of sub-optimal extreme feasible states. Each row corresponds to one futile process (*e*_2_, *e*_12_, *e*_20_, *e*_39_, *e*_40_ or *e*_41_) superimposed on the common biomass-producing growth backbone *e*_11_. Dots indicate active reactions, whereas blank entries denote inactive reactions. For active reactions, green denotes operation in the forward direction and red in the reverse direction. Reaction identifiers and their corresponding biochemical assignments are provided in Supplementary Data S2.

To visualize the structure of these candidate dissipative states, Fig. 6 reports, for each of the six associated futile processes (*e*_2_, *e*_12_, *e*_20_, *e*_39_, *e*_40_ and *e*_41_), their decomposition over the reactions of the metabolic network. Active reactions are indicated by dots, whereas inactive reactions are left blank. For active reactions, the direction of operation is also shown: green denotes the forward direction and red the reverse direction.

Across the six dissipative extreme states, the associated dissipative processes share a large invariant backbone. Among the 86 reactions represented in Fig. 6, 34 are inactive in all six processes, whereas 16 active reactions are common to all scenarios. This common active set includes the photosynthetic and respiratory electron-transport modules as well as the transhydrogenase (TH), indicating that all six processes implement a direct photosynthesis–respiration compensation mechanism.

Operationally, these dissipative processes act as energetic shunts and *electron-sink* routes that couple photochemical input to respiratory electron flow, yielding (within the dissipative process) an approximately zero net O_2_ exchange, i.e. photosynthetic O_2_ evolution is balanced by respiratory O_2_ uptake. In other words, they dissipate photochemical energy through direct photosynthesis–respiration compensation with no net contribution to biomass synthesis. The remaining 35 reactions differ across scenarios and define the state-specific routing patterns that distinguish the six dissipative motifs described below.

Independent studies have reported measurable oxidative pentose phosphate pathway (oxPPP) activity in photoautotrophic oxygenic phototrophs (Young et al., 2011). In our elementary-process analysis, the biomass-carrying growth process *e*_11_ does not recruit oxPPP. Therefore, among the dissipative candidates, the oxPPP-recruiting process *e*_20_ emerges as a particularly plausible mechanism to reconcile a conserved growth backbone with an additional light-driven energetic loss.

Process *e*_20_ corresponds to concomitant operation of the oxidative branch of the pentose phosphate pathway with the CO_2_-fixing steps of the Calvin–Benson cycle. In this motif, oxPPP short-circuits the Calvin–Benson cycle at the glucose-6-phosphate node and reinjects carbon as ribulose-5-phosphate, thereby creating a carbon-recycling loop with a net energetic cost. In the present lumped stoichiometry, this concomitant Calvin–oxPPP operation results in a net hydrolysis of one mole of ATP. Process *e*_20_ is thus energetically equivalent to process *e*_40_, in which the same net ATP dissipation is represented through the generic ATP-hydrolysis sink ATPM, but here the dissipation is implemented through a structured metabolic loop involving oxPPP and the primary carbon-fixation reactions of the Calvin–Benson cycle. Process *e*_20_ therefore provides a mechanistically more explicit route for dissipative operation under self-shading (Fig. 7).

**Figure 7:**
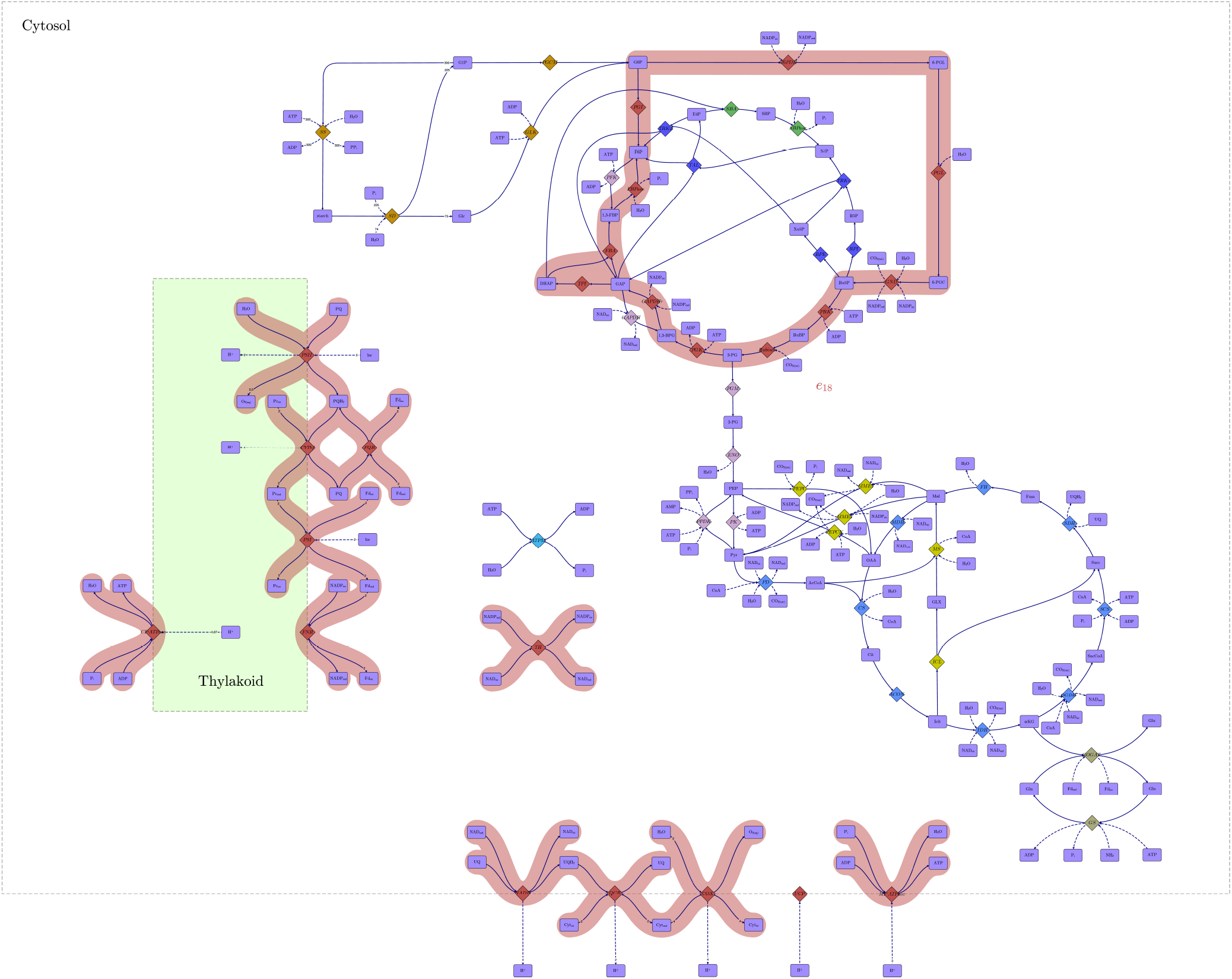
Reaction-level representation of the dissipative elementary process *e*_20_, identified as a plausible oxPPP-associated dissipative motif. The highlighted active reactions show concomitant operation of the oxidative pentose phosphate pathway and the CO_2_-fixing branch of the Calvin–Benson cycle. In this configuration, carbon is withdrawn from the Calvin cycle at the glucose-6-phosphate node and reinjected as ribulose-5-phosphate, forming a carbon-recycling loop superimposed on the photosynthetic energy-transduction backbone. In the present lumped stoichiometry, this Calvin–oxPPP loop entails a net hydrolysis of one ATP, providing a mechanistic route for dissipative operation. Reaction identifiers and their corresponding biochemical assignments are provided in Supplementary Data S2.

### Deriving an anabolic demand reaction from the growth-core process *e*_11_

Because the growth-core process *e*_11_ embeds both biomass formation and the associated energy-transduction machinery, we extracted from it an explicit anabolic demand reaction describing the metabolic requirements of biomass synthesis independently of photosynthetic and respiratory energy conversion. To this end, *e*_11_ was restricted to the subspace spanned by all reactions other than those belonging to the photosynthetic and respiratory energy-transduction modules, and the resulting reduced reaction vector was mapped through the stoichiometric matrix onto the corresponding net transformation of metabolites and cofactors (Rügen et al., 2012).

This procedure yields the following overall anabolic demand reaction, expressed per C-mole of biomass formed:

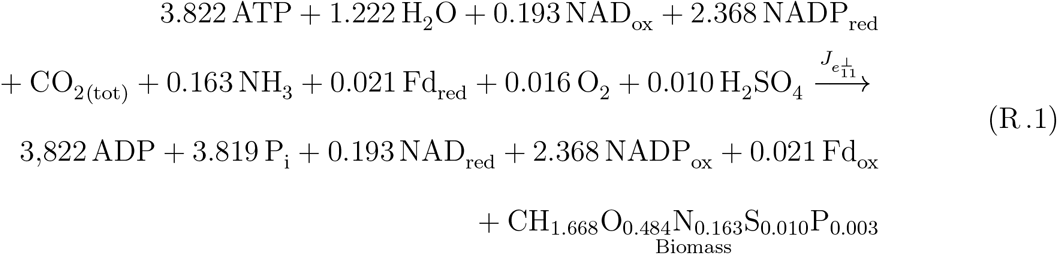

The corresponding anabolic process is characterized by two quantities that remain invariant within the present descriptor: an ATP requirement of *ν*_ATP,*x*_ = 3.822 per C-mol of biomass produced, and an effective phosphorylation-to-electron requirement *P*_2e_|_*x*_ = 1.75. Because biomass composition remains essentially unchanged, within experimental uncertainty, over the nutrient-replete growth conditions investigated here (Table 1), the corresponding anabolic demand is expected to remain nearly constant as well. These values indicate that biomass formation requires an ATP-rich energy supply relative to reducing equivalents.

This stoichiometric result provides a direct interpretation of the effective energetic coupling inferred above. Because the anabolic process embodied in the growth core requires an ATP-rich energy supply relative to reducing equivalents, the admissible photosynthesis–respiration balance is constrained by the need to satisfy this demand at steady state. In this framework, the inferred value *κ*^⋆^ suggests that photosynthetic energy conversion alone is insufficient to meet optimally the anabolic ATP requirement associated with carbon fixation. Rather, optimal photon-to-biomass conversion requires a non-zero, properly adjusted respiratory flux operating in series with photosynthesis, so as to complement the photosynthetic ATP supply and match the energetic demand of growth (Eq. (23)). This suggests that light respiration, under growth-optimal conditions, is not merely tolerated but functionally required and tightly regulated to optimize carbon assimilation. By contrast, when respiration exceeds the level required to satisfy this balance, it no longer supports anabolism but acts as an energetic drain, thereby lowering the overall efficiency of light-energy conversion into biomass.

## Discussion

### Central message

This work provides a unified, experimentally anchored reading of photon-to-biomass efficiency in an oxygenic phototroph grown in a controlled photobioreactor. First, growth aligns with a photic variable that is metabolically meaningful at the timescale of growth—the effective photochemical input *ϕ*_γ_—rather than with the incident irradiance *q*_∩_ or any local light metric. Second, the empirically observed linear scaling *µ* ∝ *ϕ*_γ_ defines a stringent constraint that selects an effective energetic partitioning within photosynthetic electron transport, summarized here by the effective photosynthetic coupling parameter *κ*^⋆^ = ATP / 2e^−^, i.e. the ATP yield per two electrons transferred. Third, once this partition is prescribed, the photosynthesis–respiration balance becomes strongly constrained by energy and redox bookkeeping (i.e. strict stoichiometric accounting of ATP and reducing equivalents), and the growth map reveals that a non-zero but properly adjusted respiration level is a condition of growth-optimal operation. In light of the ATP-rich anabolic demand extracted from the growth-core process, this suggests that respiration in the light is part of the functional bioenergetic adjustment required to optimize carbon fixation under the imposed energetic constraints (Eq. (23)). Finally, dissipative operation emerges when the photosynthesis–respiration balance shifts toward excess respiratory demand and/or when specific carbon-routing motifs become feasible, most notably Calvin–oxPPP recycling.

Taken together, these results support two central conclusions. (i) Respiration in the light is not merely a loss term: in an ATP-rich anabolic regime, it appears as a regulated support function that helps match energy and redox demand and, near the optimum, may be functionally required for maximal carbon assimilation. This is directly supported by Region I of the growth map, where insufficient respiration limits growth despite available photochemical forcing. (ii) When respiration becomes excessive relative to photosynthetic supply, it acts as an electron-sink route that lowers photon yield and pushes the system into a dissipative regime.

### Why expressing light as *ϕ*_γ_ changes the interpretation of efficiency

A central methodological choice of this study is to express the light forcing in terms of photochemically productive absorption, rather than as incident photon flux density or local irradiance. This choice is supported by the separation of time scales inherent to photobioreactors: photophysical charge-separation events occur on femto-to picosecond time scales (Mirkovic et al., 2017), mixing-induced light fluctuations occur on second time scales (Janssen et al., 2000), and growth integrates metabolic activity over hours (Takache et al., 2010). At the scale relevant to growth, cells repeatedly experience the range of irradiances present in the reactor; it is therefore natural to represent the forcing by a reactor-scale average of a local, photochemically productive absorption rate (Cornet and Dussap, 2009).

Importantly, *ϕ*_γ_ incorporates, by construction, the primary photochemical saturation through *ρ*(𝒜) (Cornet and Dussap, 2009; Takache et al., 2012), thereby distinguishing absorbed excitations that effectively drive electron transport from those that are dissipated as heat or fluorescence. When growth is expressed against *ϕ*_γ_, the near-optimal chemostat regime exhibits an approximately linear scaling with no statistically supported intercept. This does not imply that non-growth-associated demands are strictly absent. Rather, it indicates that, within this regime and given this forcing definition, an additional macroscopic “maintenance” term is not required to explain the observed growth–light relationship; such demands may be (i) comparatively small, (ii) embedded in the effective forcing definition, and/or (iii) not identifiable from the available macroscopic data.

### Bioenergetic interpretation of *κ*^⋆^ and of the anabolic demand

The inferred coupling *κ*^⋆^ should be interpreted as an effective, cell-scale energetic partitioning consistent with near-optimal operation under the imposed constraints, rather than as a universal constant of the photosynthetic machinery. In the present framework, *κ*^⋆^ summarizes how photosynthetic electron transport delivers ATP relative to reductant supply under growth-optimal operation, thereby setting the bioenergetic leverage available to satisfy anabolism.

The anabolic-demand analysis extracted from the growth-core process *e*_11_ provides a mechanistic rationale for why this leverage matters. Because the biomass pseudo-reaction is parameterized from experimentally measured macromolecular composition of *Chlamydomonas* (Table 1), which varies only within measurement uncertainty across the investigated chemostat conditions, the resulting anabolic demand can be viewed as a regime-level requirement rather than an artifact of a particular operating point. In the reduced descriptor, biomass formation is characterized by (i) an ATP requirement per unit biomass produced, *ν*_ATP,*x*_ = 3.822, and (ii) an effective electron-equivalent requirement summarized here by *P*_2e_|_*x*_ = 1.75 (an effective phosphorylation-to-electron requirement for biomass formation). Together, these quantities indicate that biomass formation is ATP-rich relative to reductant demand. Consistently, satisfying such a demand at steady state constrains the admissible ATP return per photosynthetic electron (captured here by *κ* = ATP / 2e^−^). In this framework, the inferred value *κ*^⋆^ suggests that photosynthetic energy conversion alone does not meet optimally the ATP requirement associated with carbon fixation and biomass synthesis. Rather, a non-zero and properly adjusted respiratory flux becomes functionally necessary for growth-optimal operation, because respiration acting in series with photosynthesis provides the additional energetic flexibility required to match the ATP-rich anabolic demand. This, in turn, suggests that light respiration under near-optimal growth conditions should not be viewed primarily as a passive loss term, but as a regulated support function contributing to ATP–redox matching and optimal carbon assimilation. Conversely, once respiratory activity exceeds the level required for this support role, it becomes a net energetic drain that increases the photochemical demand for a fixed growth outcome.

Encouragingly, the magnitude of *κ*^⋆^ is also consistent, within uncertainties, with independent stoichiometric/biophysical estimates of the ATP return of the photosynthetic chain in *Chlamydomonas*. Using a chloroplast ATP-synthase coupling of 14 H^+^ per 3 ATP (i.e. 14*/*3) and the usual proton-translocation bookkeeping for linear electron flow, Alric et al. (2010, 2014) show that linear photochemistry alone provides insufficient ATP for carbon fixation and that a modest cyclic electron flow (CEF) is required to close the ATP budget. In particular, the required CEF is estimated to be on the order of ∼ 6–16% of the linear electron flow under aerobic conditions (Alric et al., 2010), a range that increases the effective ATP return from ≃ 1.29 (linear only) to ≃ 1.34–1.42 ATP per 2 e^−^. This is consistent, within uncertainties, with our inferred *κ*^⋆^ ≈ 1.45. Both estimates are subject to uncertainty, as they rely on effective proton/ATP stoichiometries and on simplified partitioning assumptions for CEF and auxiliary proton/electron pathways. This supports interpreting *κ*^⋆^ as a realistic operating-point descriptor reflecting ATP–redox rebalancing through CEF (and associated regulation) under growth-optimal conditions.

Finally, the effective coupling between photochemical electron transfer and ATP synthesis can also be examined through thermodynamics of irreversible processes (TIP), which relates coupling, driving forces and dissipation in energy-conversion modules and has been discussed in the context of photobioreactors (Cornet et al., 1998). In this view, ratios of the ATP / 2e^−^ type are naturally interpreted as operating-point descriptors that emerge from the balance between productive conversion and unavoidable dissipation, rather than as fixed biophysical constants. While we do not pursue a TIP-based derivation here, the inferred *κ*^⋆^ offers a concrete quantitative target for such complementary analyses and for future dissipation-aware coupling studies.

### The two-dimensional growth map: regimes and an operational optimality rule

A key contribution of this work is the two-dimensional growth map in the (*q*_γ_, *J*_COX_) plane, constructed under fixed bioenergetic parametrization. This map exposes two qualitatively distinct regimes. In Region I, growth is respiration-limited: increasing respiratory flux increases growth potential at nearly fixed photochemical input, indicating that respiratory electron-processing capacity (redox balancing and electron-sink accommodation) constrains growth. This regime provides a mechanistic underpinning for the idea that respiration in the light can support efficiency rather than merely reduce it (Salinas-Giégé et al., 2014; Kaye et al., 2019). In Region II, growth becomes dissipative (over-respiring): maintaining a given growth rate requires increasing photochemical input as respiratory activity increases, implying a loss of photon yield.

Separating both regions is a locus of maximal growth potential (the maximal-growth ridge), along which the photosynthesis–respiration partition follows an approximately constant ratio in O_2_ units (Eq. (23)). This relationship can be interpreted as an operational optimality rule: for each available photochemical input, there exists an optimal respiration level that maximizes growth under stoichiometric, thermodynamic, and bioenergetic constraints. This complements classical photobioreactor interpretations of kinetic versus radiative-transport-limited operation (e.g. via the illuminated fraction or the onset of photolimitation) by providing a mechanistic description of how intracellular energy/redox balancing selects an optimal photosynthesis–respiration balance at the reactor scale (Cornet et al., 1998; Cornet and Dussap, 2009; Takache et al., 2010).

### Dissipative mechanisms: why Calvin–oxPPP recycling is a strong candidate

Moving beyond regime classification requires linking the macroscopic signature of dissipation to intracellular routing patterns that can realize it under constraints. Here, extreme-state enumeration, thermodynamic screening, and elementary-process decomposition yielded a short list of dissipative scenarios compatible with the operationally sub-optimal steady state. Across these scenarios, the dissipative processes share a common backbone that includes photosynthetic and respiratory electron transport and the transhydrogenase. This points to a generic energetic-shunt architecture: photochemical input is diverted toward O_2_-reducing electron-sink routes, yielding (within the dissipative process itself) an approximately zero net O_2_ exchange, i.e. photosynthetic O_2_ evolution is locally compensated by respiratory O_2_ uptake.

Such light-dependent engagement of electron sinks is not specific to eukaryotic microalgae. In cyanobacteria, thylakoid terminal oxidases and other auxiliary outlets can contribute to O_2_ uptake in the light and act as controllable sinks when photosynthetic electron supply and downstream metabolic demand become mismatched, providing a conceptually related route to dissipative operation in oxygenic phototrophs (Ermakova et al., 2016).

Among the candidate motifs emerging from our analysis, the Calvin–oxPPP scenario stands out as both parsimonious and consistent with prior evidence. A growing body of work has highlighted that partial operation of oxPPP in the light can occur via the “glucose-6-phosphate shunt” around the Calvin–Benson cycle (Sharkey and Weise, 2016), and flux-analysis evidence under photoautotrophy supports non-negligible oxPPP activity in oxygenic phototrophs (Young et al., 2011; Cahoreau, 2012). In our elementary-process decomposition, the growth-carrying process *e*_11_ does not recruit oxPPP, whereas the dissipative candidate *e*_20_ does. This makes *e*_20_ a natural mechanism to reconcile a conserved growth module with an additional light-driven energetic loss.

Mechanistically, *e*_20_ corresponds to concomitant operation of the oxidative branch of the pentose phosphate pathway with the CO_2_-fixing steps of the Calvin–Benson cycle. OxPPP short-circuits the Calvin cycle at the glucose-6-phosphate node and reinjects carbon as ribulose-5-phosphate, thereby creating a carbon-recycling loop with a net energetic cost; in the present lumped stoichiometry, this cost appears as a net hydrolysis of one mole of ATP. The plausibility of such Calvin–oxPPP concurrency is reinforced by the tight light/redox regulation of the oxPPP entry enzyme glucose-6-phosphate dehydrogenase (G6PDH), which is down-regulated in the light by thioredoxin and can be reactivated upon transitions to the dark or oxidizing conditions (Née et al., 2009). Together, these observations provide a concrete route by which self-shading and light-history constraints can gate a dissipative mode without altering the structural biomass-demand module itself.

### Implications for monitoring and control

A practical implication of the present framework is that near-optimal operation can be formulated as a control problem in which the photosynthesis–respiration balance is regulated to remain close to the maximal-growth ridge. In this view, respiration is necessary but must be tuned relative to photosynthetic activity: too little respiration constrains growth in the respiration-limited regime (Region I), whereas excessive respiration pushes the system into a dissipative regime (Region II) and lowers photon yield. The ridge relationship (Eq. (23)) therefore provides an operational setpoint linking respiratory demand to photosynthetic supply (Bernard and Rémond, 2012; Martínez et al., 2018).

This perspective suggests that improved monitoring of respiratory activity in the light— for instance via indirect reconstruction from online gas-exchange measurements combined with model-based constraints—could enable feedback strategies that maintain cultures near their optimal balance despite slow drifts in biomass concentration, self-shading, and lightexposure patterns (Cogne et al., 2025; Eriksen et al., 2007). More broadly, the framework provides a quantitative bridge between measurable reactor-scale signals (effective photochemical input, gas exchange) and intracellular energetic motifs, thereby supporting both mechanistic interpretation and process-oriented optimization.

### Limitations and generalizability

The proposed framework is intentionally parsimonious. The metabolic descriptor preserves the energy/redox accounting required to constrain feasible photosynthesis–respiration balances and photon yield, but it does not resolve compartment-specific metabolite pools, transporter-level regulation, or fast acclimation dynamics. Accordingly, the identified dissipative motifs should be interpreted as feasible pathway-level routes consistent with the imposed constraints, rather than as uniquely realized *in vivo*.

Several parameters were fixed to define a parsimonious baseline (e.g. respiratory P/O yields and the photochemical saturation law *ρ*(𝒜)). While these assumptions are standard for the investigated regime, their quantitative values may shift with physiology, strain, and reactor conditions; future work should evaluate how sensitive the ridge structure and motif ranking are to such variations and to explicit representations of uncoupling and alternative electron sinks.

Finally, although the numerical values reported here are specific to *Chlamydomonas reinhardtii* and to the present photobioreactor setting, the conceptual structure of the framework—expressing forcing as photochemically productive absorption, inferring an effective chain-level energetic partition, and mapping feasible growth across photosynthesis–respiration space—should also be relevant for other oxygenic phototrophs in which electron-sink routes and carbon-recycling loops contribute to photon-to-biomass efficiency. In that respect, available oxygen-exchange measurements reported for cyanobacteria under dilute and well-mixed conditions with limited light gradients suggest respiratory oxygen-uptake levels of only a few percent of gross photosynthetic oxygen evolution, i.e. in the same order of magnitude as the ratio predicted here by Eq. (23) (Ermakova et al., 2016). Such a comparison must nevertheless remain cautious, since in cyanobacteria the respiratory and photosynthetic electron-transfer chains are partly intertwined within the same thylakoid membrane system and share common redox components, unlike in eukaryotic microalgae such as *Chlamydomonas reinhardtii*.

## Conclusions

In this study, we combined a quantified description of the radiative environment with a thermodynamically constrained metabolic framework to relate light history, bioenergetic coupling, and photon-to-biomass efficiency in an oxygenic phototroph.

First, we showed that growth in the near-optimal chemostat regime is captured by an experimentally grounded invariant when light is expressed as an effective photochemical input *ϕ*_γ_, i.e. the reactor-scale average rate of photochemically productive absorption. This reformulation provides a physiologically relevant forcing at the timescale of growth and defines an operationally constant biomass yield on productive photons in the explored regime.

Second, imposing this invariant enabled inference of an effective photosynthetic energetic coupling, *κ*^⋆^ = ATP / 2e^−^, interpreted as a cell-scale partition of photochemical energy that is consistent with growth-optimal operation under stoichiometric and thermodynamic constraints. Fixing this coupling strongly constrains the admissible photosynthesis–respiration balance through energy and redox bookkeeping.

Third, the resulting two-dimensional growth map in the (*q*_γ_, *J*_COX_) plane revealed a respiration-limited region and a dissipative (over-respiring) regime, separated by a locus of maximal growth potential. Along this ridge, optimal operation is associated with a characteristic balance between respiratory demand and PSII activity (Eq. (23)), showing that a non-zero but properly adjusted respiration level is required to maximize photon-to-biomass conversion, whereas excessive respiration acts as an electron-sink route that lowers efficiency.

Finally, systematic exploration of the feasible flux space at an operationally sub-optimal (self-shaded) state, including vertex enumeration, thermodynamic filtering, and extreme-ray decomposition, identified a small set of candidate dissipative motifs. Among them, concomitant operation of the Calvin–Benson cycle and the oxidative pentose phosphate pathway (oxPPP) emerged as a parsimonious mechanism for dissipative operation, implementing a carbon-recycling loop with a net energetic cost while preserving the growth-driving backbone.

Beyond providing mechanistic insight, the framework also suggests a process-oriented implication: maintaining cultures near the maximal-growth ridge requires control of the photosynthesis–respiration balance. Methods to monitor or reconstruct respiratory activity at reactor scale (e.g. through gas-exchange signals combined with model-based estimation) could therefore enable feedback strategies that keep cultures near their optimal energetic state despite drifts in self-shading and light-exposure patterns.

## Acknowledgments

The authors gratefully acknowledge Hosni Takache, whose doctoral work and experimental results constitute a major foundation of the present study, although he was not directly supervised by the corresponding author.

The authors also gratefully acknowledge Arnaud Martzolff for his substantial experimental contribution to this work, notably through the generation and analysis of the data underlying the present study.

A significant part of this work was supported by the French National Research Agency through the ANR projects ALGOMICS (ANR-08-BIOE-002) and ChloroPaths (ANR-14-CE05-0041).

The authors also sincerely thank Prof. Dr. Alexander Bockmayr for the opportunity to receive training in constraint-based and computational approaches for metabolic network analysis through the co-supervision of two Master’s theses, which played an important role in shaping the methodological framework developed in this study.

## Data availability statement

The data supporting the results of this study are included in the article and its Supplementary Data. Additional materials and intermediate analysis files are available from the corresponding author upon reasonable request.

## Appendices

### A A second-law bound for the maximal primary photochemical yield *ρ*_max_

This Appendix derives a second-law upper bound on the primary photochemical yield *ρ*(𝒜) from the exergy content of radiation. Because *ρ*(𝒜) reaches its maximum *ρ*_max_ in the lowexcitation limit (𝒜 ≪ *K*) of Eq. (4), the bound provides a physically motivated ceiling for *ρ*_max_, and therefore constrains the effective photochemical input *ϕ*_γ_ defined in Eq. (6). The derivation below combines a standard radiative-exergy argument with an isotropic photon-gas closure and an effective-medium correction of the pressure-work term; the resulting expression should therefore be interpreted as an effective upper-bound estimate rather than as a universal closed-form identity.

#### A.1 Radiative exergy and convertible fraction

Consider a control volume at temperature *T* receiving radiation from a source represented, for bounding purposes, by an effective emission temperature *T*_*S*_. Let 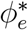 denote the incoming radiative energy flux per unit volume of reactive system. By the second law, only part of this incoming radiative power can be converted into work-like free energy; the remainder is necessarily degraded into entropy at temperature *T*.

Following standard exergy-of-radiation arguments (Petela, 1964; Jeter, 1981; Landsberg and Tonge, 1980; Würfel, 1982), the maximal convertible part of the incoming radiation combines an energy term, a mechanical work contribution arising from radiation pressure, and an entropic penalty. In the present setting, however, the pressure-work contribution must be evaluated consistently with propagation in a refractive medium.

For an isotropic radiation field (photon-gas closure), the radiation pressure satisfies

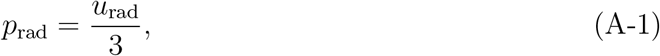

so that the corresponding mechanical contribution scales as

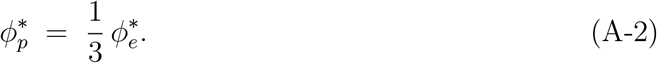

To account, within the present effective-medium description, for the modified relation between radiative energy transport and momentum transfer in a refractive medium, we introduce the phenomenological scaling

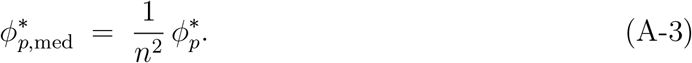

where *n* denotes the refractive index of the medium.

We then define the maximal convertible radiative contribution as

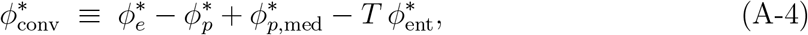

where 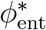 is the entropy flux associated with the incoming radiation. For isotropic blackbody radiation emitted at temperature *T*_*S*_,

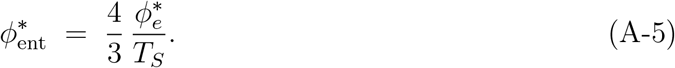

Substituting Eqs. (A-2), (A-3) and (A-5) into Eq. (A-4) yields

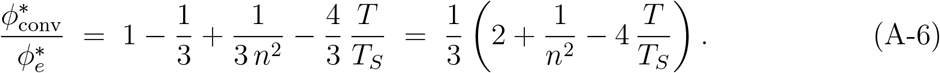

Equation (A-6) should therefore be interpreted within the present isotropic photon-gas and effective-medium framework.

#### A.2 Bound on *ρ*_max_

In our model, *ρ*(𝒜) denotes the primary quantum yield, i.e. the fraction of absorbed photons effectively generating reaction-center charge separation. In Eq. (4), this yield reaches its maximal value *ρ*_max_ in the low-excitation limit 𝒜 ≪ *K*.

A necessary upper-bound condition is that the rate of free-energy storage associated with primary photochemistry cannot exceed the maximal convertible part of the incoming radiative power. Denoting by 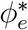 the incoming radiative energy flux, this gives the general inequality

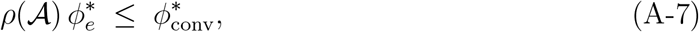

and therefore

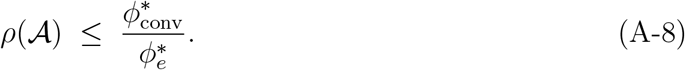

In the low-excitation regime, where *ρ*(𝒜) = *ρ*_max_, this directly yields

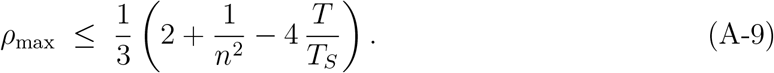

For aqueous photobioreactors, *n* ≃ 1.33 is commonly used. Taking representative ambient conditions *T* = 298 K and a solar benchmark *T*_*S*_ = 6000 K gives

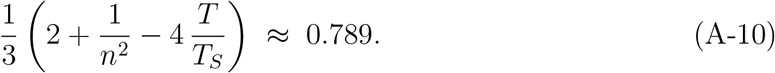

This provides a convenient benchmark close to 0.79, and supports using *ρ*_max_ = 0.8 in the main text as a physically grounded upper bound for the low-excitation primary yield. Within the present modeling framework, this corresponds to the regime in which primary photochemical conversion is assumed to operate close to its maximal efficiency.

#### A.3 Consistency check for LED illumination

The parameter *T*_*S*_ has a direct thermodynamic meaning for thermal radiation, whereas laboratory LED illumination is non-thermal. As a consistency check, we therefore define an effective temperature 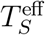 as the blackbody temperature whose mean photon energy over the PAR window (400–700 nm) matches that of the measured LED spectrum.

Let *q*_*λ*_ denote the measured spectral photon flux density. The mean LED photon energy is

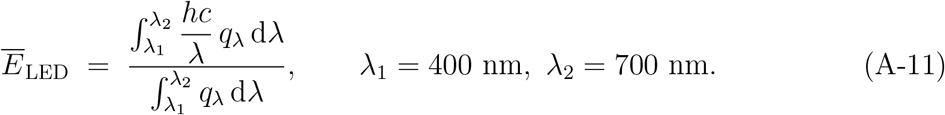

For a blackbody at temperature *T*_*S*_, the photon-number spectral density in wavelength can be written, up to a multiplicative constant that cancels in the ratio, as

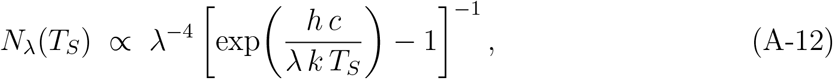

which gives

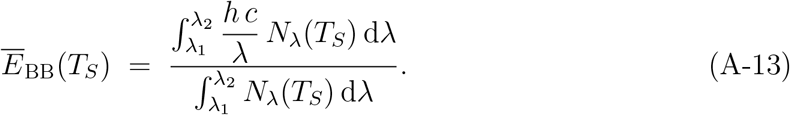

We then define 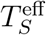 by 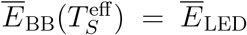. For the present LED spectrum, this yields 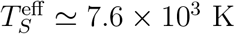.

Using *T* = 298 K, *n* = 1.33, and 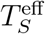 in Eq. (A-9) gives again *ρ*_max_ ≲ 0.80, i.e. very close to the solar-benchmark value.

### B Explicit MILP form of thermodynamic feasibility constraints

This appendix gives the explicit mixed-integer linear formulation used to enforce thermodynamic feasibility by coupling reaction flux directions to reaction driving forces (affinities).

#### B.1 Variables and driving forces

Let ***ν*** ∈ ℝ^*m*×*r*^ be the stoichiometric matrix and **J** ∈ ℝ^*r*^ the vector of intracellular reaction fluxes (biomass-specific rates). Introduce **x** = ln **c** ∈ ℝ^*m*^, where **c** denotes intracellular concentrations, with prescribed bounds

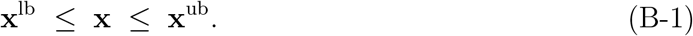

Chemical potentials and affinities are written as

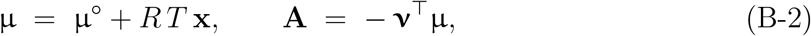

so that **A** is affine in **x**. This concentration-based formulation assumes ideal-solution behavior (unit activity coefficients), with water activity set to unity.

#### B.2 Three-state directional logic and MILP constraints

For each reaction *i*, we enforce a three-state thermodynamic logic of De Donder type: (i) forward operation (*J*_*i*_ ≥ 0 and *A*_*i*_ ≥ *ϵ*), (ii) reverse operation (*J*_*i*_ ≤ 0 and *A*_*i*_ ≤ −*ϵ*), or no net flux at equilibrium (*J*_*i*_ = 0 and *A*_*i*_ = 0). To encode this logic, we introduce two binary variables 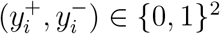 such that

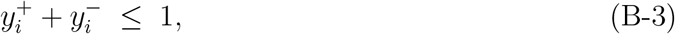

where 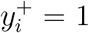 selects the forward mode and 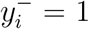 selects the reverse mode. Let *ϵ >* 0 be a small tolerance used to replace strict inequalities and to separate active forward/reverse states from the equilibrium state. The logic above is enforced by the following linear inequalities:

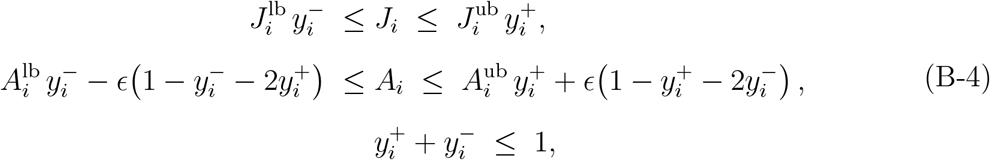

where 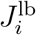 and 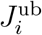 denote the lower and upper bounds on *J*_*i*_, and 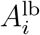 and 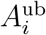 are bounds on *A*_*i*_ derived from concentration bounds (Section B.3). Together, Eq. (B-4) enforces sign consistency between *J*_*i*_ and *A*_*i*_ for non-zero fluxes, while allowing the equilibrium/zero-flux state when 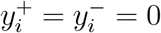.

##### Loop thermodynamic consistency

In addition to the reaction-wise sign constraints of Eq. (B-4), thermodynamic feasibility requires consistency over internal stoichiometrically closed loops. This is enforced through the linear equality

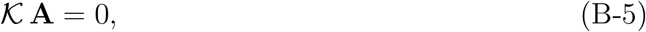

where 𝒦 is the null-space basis associated with the internal stoichiometric matrix. The condition states that the sum of affinities over any internal stoichiometrically closed loop must vanish.

#### B.3 Bounds on affinities

Affinity bounds are obtained from the log-concentration bounds **x**^lb^ ≤ **x** ≤ **x**^ub^ (Eq. (B-1)) and the affine dependence **A**(**x**) (Eq. (B-2)). For each reaction *i*, conservative bounds 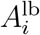 and 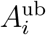 are obtained by propagating the interval **x** ∈ [**x**^lb^, **x**^ub^] through the affine expression of *A*_*i*_(**x**) (e.g. by interval arithmetic), or equivalently by evaluating *A*_*i*_ at the vertices of the box and taking the minimum/maximum values.

### C Linear energetic-coupling constraints used in selected analyses

For the analyses in which effective energetic coupling was prescribed, the following linear equality constraints were added to the stoichiometric model.

For photosynthesis:

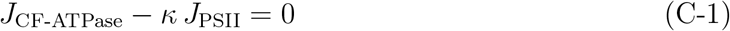

which fixes the effective ATP yield per two electrons transferred through the photosynthetic module, denoted *κ*.

For respiration:

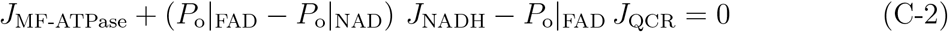

which enforces stoichiometric consistency between mitochondrial ATP synthesis and respiratory electron transfer under prescribed NADH- and FAD-linked P/O ratios.

Here, *J*_CF-ATPase_ and *J*_MF-ATPase_ denote the chloroplastic and mitochondrial ATP synthases, respectively; *J*_PSII_ denotes PSII water oxidation; *J*_NADH_ the NADH dehydrogenase flux; and *J*_QCR_ the ubiquinone–cytochrome *c* reductase flux.

