## Supplementary data S1 for "A mechanistic framework for photon-to-carbon efficiency: coupling photosynthetic energy supply and light respiration in microalgae"

### Input data and spectral computation used for the evaluation of $\phi_\gamma$

This Supplementary Data documents the spectral computation used to evaluate the effective photochemical input $\phi_\gamma$, together with the corresponding input data used for the reconstruction of the wavelength-dependent radiative properties and for the computation of the radiative field in the culture.

### S1.1. General principle

In the main text, the effective photochemical input is introduced in reduced form as the depth-averaged rate of photochemically productive absorption,

$$\phi_\gamma \;=\; \frac{1}{L}\int_0^L \rho(\mathcal{A}(z))\;\mathcal{A}(z)\,\mathrm{d}z, \tag{1}$$

where $L$ is the illuminated reactor depth, $\mathcal{A}(z)$ is the local biomass-normalized absorption (antenna excitation) rate, and $\rho(\mathcal{A})$ is the primary photochemical yield.

In practice, $\phi_\gamma$ was evaluated spectrally over the PAR domain by combining: (i) the measured incident spectral photon flux density, (ii) the wavelength-dependent radiative properties of the culture, and (iii) the local photochemical conversion law.

### S1.2. Input data used for optical-property reconstruction

The reconstruction of the wavelength-dependent radiative properties follows the inverse optical-property framework of Dauchet et al. [2015]. For *Chlamydomonas reinhardtii*, this reconstruction relies on the geometric and pigment-related descriptors previously established in Takache et al. [2010, 2012].

In the range of conditions investigated here, the cell-shape model, the cell-size distribution, and the relative pigment distribution were assumed unchanged, consistently with the limited variability reported in those studies. By contrast, the total pigment content was specified for each analyzed steady state and used in the reconstruction of the spectral radiative properties.

Biomass concentration does not enter the reconstruction of the intrinsic radiative properties themselves. It is used only subsequently in the computation of the radiative field $G_\lambda(z)$ within the culture and therefore in the evaluation of $\mathcal{A}(z)$ and $\phi_\gamma$.

#### S1.2.1. Common inputs assumed for all analyzed states

Table S1a summarizes the geometric descriptors and relative pigment distribution used as common inputs for the optical-property reconstruction.

**Table S1a.** Common inputs used for optical-property reconstruction.

| Quantity | Symbol | Value | Unit | Source / comment |
|---|---|---|---|---|
| Cell-shape model | – | Tchebychev particle T2(0.084), order 2 | – | Takache et al. [2012] |
| Deformation parameter | $\varepsilon$ | 0.084 | – | Takache et al. [2012] |
| Size-distribution type | – | log-normal | – | Takache et al. [2012] |
| Equivalent radius | $r_{\mathrm{eq}}$ | 4.0 | µm | Takache et al. [2010, 2012] |
| Size-distribution parameter | $\ln\sigma$ | 0.17 | – | Takache et al. [2012] |
| Relative chlorophyll a fraction | – | 59.0 | % | Takache et al. [2012] |
| Relative chlorophyll b fraction | – | 23.5 | % | Takache et al. [2012] |
| Relative PPC fraction | – | 17.5 | % | Takache et al. [2012] |

#### S1.2.2. State-specific inputs

For each analyzed steady state, the total pigment content was used in the reconstruction of the spectral radiative properties, while the biomass concentration $C_x$ was used in the computation of the radiative field and of the corresponding effective photochemical input.

For the additional steady states analyzed in the present study (States 7 and 8), all state-specific input data used in the spectral computation, including dilution rate, biomass concentration, and total pigment content, were taken from Martzolff [2013].

Table S1b summarizes these state-specific inputs.

**Table S1b.** State-specific inputs used for spectral optical-property reconstruction and radiative-field computation.

| State ID | $q_{\cap}$ $(\mu\mathrm{mol}_{\mathrm{h}\nu}\cdot\mathrm{m}^{-2}\cdot\mathrm{s}^{-1})$ | $D$ $(\mathrm{h}^{-1})$ | $w_{\mathrm{pig}}$ (% w/w) | $C_x$ $(\mathrm{kg}\cdot\mathrm{m}^{-3})$ |
|---|---|---|---|---|
| State 1 | 110 | 0.032 | 3.50 | 0.35 |
| State 2 | 200 | 0.041 | 3.08 | 0.41 |
| State 3 | 300 | 0.054 | 2.84 | 0.47 |
| State 4 | 400 | 0.057 | 2.66 | 0.51 |
| State 5 | 500 | 0.058 | 2.54 | 0.57 |
| State 6 | 600 | 0.061 | 2.44 | 0.60 |
| State 7 | 200 | 0.041 | 3.05 | 0.42 |
| State 8 | 200 | 0.016 | 5.37 | 0.59 |

States 1–6 correspond to the chemostat dataset reported in Takache et al. [2012]. States 7 and 8 correspond to the additional steady states reported in Martzolff [2013].

### S1.3. Incident light spectrum

Let $q_{\cap,\lambda}$ denote the incident spectral photon flux density ($\mathrm{mol_{h\nu}} \cdot \mathrm{m}^{-2} \cdot \mathrm{s}^{-1} \cdot \mathrm{nm}^{-1}$), measured over the PAR window

$$\lambda \in [400, 700] \text{ nm}.$$

The normalized incident light spectrum used in the spectral computation is provided as a text file together with this Supplementary Data.

### S1.4. Spectral radiative properties

For each experimental condition, wavelength-dependent radiative properties were described by:

- the mass absorption coefficient $E_{a,\lambda}$,
- the mass scattering coefficient $E_{s,\lambda}$,
- and the backscattered fraction $b_\lambda$.

These spectral optical properties were retrieved by applying the inverse optical-property identification method of Dauchet et al. [2015] to the *Chlamydomonas reinhardtii* chemostat dataset of Takache et al. [2010, 2012] and to the additional steady states analyzed in the present work.

### S1.5. Spectral radiative field

At each wavelength, the local irradiance field $G_\lambda(z)$ was reconstructed using the same two-flux formalism as in the main text, applied wavelength by wavelength. In the one-dimensional planar geometry of the reactor, this yields a spectral irradiance profile of the form

$$\frac{G_\lambda(z)}{q_{\cap,\lambda}} \approx \exp\left[-\frac{1+\alpha_\lambda}{2\,\alpha_\lambda}\, E_{a,\lambda}\, C_x\, z\right], \tag{2}$$

with

$$\alpha_\lambda = \sqrt{\frac{E_{a,\lambda}}{E_{a,\lambda} + 2\, b_\lambda\, E_{s,\lambda}}}. \tag{3}$$

Here, $C_x$ denotes the biomass concentration and $z$ the depth coordinate measured from the illuminated surface.

### S1.6. Spectral absorption and effective excitation rate

The wavelength-resolved biomass-normalized absorption rate was defined as

$$\mathcal{A}_\lambda(z) = E_{a,\lambda}\, G_\lambda(z). \tag{4}$$

The total local antenna excitation rate was then obtained by spectral integration over the PAR range:

$$\mathcal{A}(z) = \int_{400}^{700} \mathcal{A}_\lambda(z)\,\mathrm{d}\lambda. \tag{5}$$

The primary photochemical yield was evaluated from this total local excitation rate according to the same hyperbolic law as in the main text,

$$\rho(\mathcal{A}) = \rho_{\max}\frac{K}{K+\mathcal{A}}. \tag{6}$$

Accordingly, the local rate of photochemically productive absorption was written as

$$\varphi_\gamma(z) = \rho(\mathcal{A}(z)) \int_{400}^{700} \mathcal{A}_\lambda(z)\,\mathrm{d}\lambda = \rho(\mathcal{A}(z))\,\mathcal{A}(z). \tag{7}$$

### S1.7. Reactor-scale effective photochemical input

The effective photochemical input finally used in the manuscript was obtained by averaging this local photochemically productive absorption rate over the illuminated depth:

$$\phi_\gamma = \frac{1}{L}\int_0^L \varphi_\gamma(z)\,\mathrm{d}z = \frac{1}{L}\int_0^L \rho(\mathcal{A}(z))\,\mathcal{A}(z)\,\mathrm{d}z. \tag{8}$$

Thus, the reduced formulation reported in the main text is the compact depth-integrated form of a spectral calculation in which the local excitation rate $\mathcal{A}(z)$ itself results from prior integration of wavelength-resolved absorption over the PAR domain.

### S1.8. Reduced wavelength-averaged formulation used in the main text

For readability, the main manuscript reports a reduced formulation written as if the optical parameters $E_a$, $E_s$, and $b$ were wavelength-independent effective quantities. This reduced formulation leads to the wavelength-averaged expressions reported in the main text and preserves the same conceptual structure: radiative attenuation $\rightarrow$ local absorption $\rightarrow$ primary photochemical conversion $\rightarrow$ reactor-scale averaging.

The full spectral computation was used for numerical evaluation of $\phi_\gamma$, whereas the reduced formulation was retained in the paper to keep the presentation analytically transparent.

### S1.9. Data provided

This Supplementary Data includes:

- the normalized incident light spectrum over the PAR range, provided as a text file;

- the geometric and pigment-related inputs used for optical-property reconstruction, including the assumed cell-shape model, the size-distribution parameters, and the relative pigment distribution;
- the total pigment contents specified for the analyzed steady states;
- the biomass concentrations $C_x$ used for the computation of the radiative field for each analyzed steady state;
- the input data used to reconstruct the wavelength-dependent radiative properties in the spectral computation, following Takache et al. [2010, 2012] and the inverse optical framework of Dauchet et al. [2015];
- and the numerical implementation used to reconstruct $G_\lambda(z)$, $\mathcal{A}(z)$, and $\phi_\gamma$.
